# SARM1 is required for macrophage immunophenotype switching that is essential for nerve repair

**DOI:** 10.64898/2026.04.07.716973

**Authors:** Julianna Bennett, Halimah Adesunkanmi, Noah Leever, Grace Bergeron, Josh Small, Cathryn Holladay, Grace Saxman, Rachel E. Williamson, Manshi Swain, Gavin Pearson, Manav Patel, Ashley L. Kalinski

**Affiliations:** Department of Biology, Ball State University, Muncie IN 47306, USA; Department of Biological Sciences, University of South Carolina, Columbia SC 29208, USA; SmartState Center for Childhood Neurotherapeutics; University of South Carolina, Columbia, SC 29204 USA

**Keywords:** Macrophages, Inflammation, Regeneration, SARM1, Wallerian Degeneration

## Abstract

SARM1 is a key executor of Wallerian degeneration in axons. Global knockout of *sarm1* in mice delays degeneration for several weeks. Recently, we reported that Schwann cell reprogramming, inflammation, and axon regeneration are also delayed in these animals. Several studies have also indicated that SARM1 has essential regulatory functions in macrophages (Mɸ). However, the role of SARM1 in Mɸ in the context of peripheral nerve injury remains unknown. Here, we report that loss of *sarm1* impairs splenic Mɸ from adopting immunological stimuli driven immunophenotypes in culture. Through a combination of cell culture, Western blotting, gene expression analysis, *in vivo* injection of Mɸ into sciatic nerves, and generation of cell specific *sarm1* conditional knockout mouse lines, we found that SARM1 is required for proper immunophenotypes in Mɸ. Loss of *sarm1* in macrophages increases neurite length of sensory neurons in culture but delays regeneration in a model of peripheral nerve injury. We identified dysregulation of several inflammatory and anti-inflammatory immunological stimuli pathways and altered regulation of both iNOS and Arginase-1 in *Sarm1-/-* Mɸ. In culture, *Sarm1-/-* Mɸ display difficulty phagocytosing and clearing myelin debris and this was recapitulated *in vivo* with a Mɸ specific *sarm1* knockout line. Generation of Mɸ and neuronal *sarm1* conditional knockout mice further indicated that SARM1 is required in both cell types for an efficient response to peripheral nerve injury. This study provides the first evidence that SARM1 signaling in Mɸ is required for injury induced inflammation, degeneration, and axon regeneration.

## Background

Axons of the peripheral nervous system (PNS) can spontaneously regenerate following traumatic injury. However, this is not the case for axons of the central nervous system (CNS) which axons fail to regenerate without intervention. This dichotomy can be attributed to differences in both neuronal intrinsic growth capacities and extrinsic environments. It has been well documented that, while CNS neurons usually fail to regenerate following an injury, they are intrinsically capable of initiating regeneration programs when their environment is altered to become more permissive^1^. In the PNS, the extrinsic environment is supportive of regeneration due to an abundance of supportive resident and infiltrating non-neuronal cells ^2^. While Schwann cells are known to support regeneration in the PNS, recent focus has been on the role of injury activated macrophages (Mɸ)^3^.

Mɸ are derived from myeloid progenitor cells in the bone marrow and become resident in tissues during development. These tissue resident Mɸ are long lived and are important for initiating responses to infection, stress, and injury^4^. These cells are replenished from monocyte precursors and will finish maturation in the corresponding tissue, where they have the ability to self-renew. Mɸ are, as a whole, extremely plastic and heterogeneous. They are susceptible to their environment and can be polarized towards more pro- or anti-inflammatory immunophenotype. While Mɸ are classically denoted M1 versus M2 Mɸ, respectively, we now know that this classification is not a clear divide and most Mɸ can display both M1 and M2 phenotypes simultaneously. It is clear however, that these Mɸ subtypes play different roles during infection, stress, and injury^4^.

Single-cell RNA profiling studies of the uninjured rodent sciatic nerve have revealed a cellular landscape composed of immune cells, mesenchymal cells, fibroblasts, Schwann cells, endothelial cells and other vascular cells^5^. Mɸ are by far the largest immune cell population in the nerve followed by monocyte-derived dendritic cells and mast cells. Following injury, the cellular landscape drastically shifts and is marked by an accumulation of bone-marrow derived myeloid cells^5^. Granulocytes and monocytes are recruited to the injury site and distal nerve stump. Granulocytes are short lived, peaking in the nerve at 1 day post-injury and dropping in significant numbers by 3 days^5^. The monocyte/Mɸ population peaks around 3 days following injury and slowly decreases at 7 days. Recent studies have shown that the recruited blood-borne monocytes mature into Mɸ during this window^2^. Several groups, including ours, have established that Mɸ are important for phagocytosis, efferocytosis, and Schwann cell reprogramming after injury^2,6^. These roles are especially important during Wallerian degeneration, a programmed destruction that leads to the removal of distal axons and debris after 5-7 days post-injury and occurs very efficiently in the PNS. SARM1 (sterile alpha and TIR motif containing 1) is a toll-like adaptor protein with NADase enzymatic function responsible for the critical depletion of NAD, effectively kickstarting Wallerian degeneration. Germline deletion of *sarm1* in rodents leads to the delay of Wallerian degeneration by approximately 2 weeks^7,8^. While it is clear that the NADase properties of SARM1 within axons are important for triggering and/or sustaining the Wallerian degeneration process, recent work has suggested that SARM1 plays additional roles in the injury response^6,9^. We have recently reported that following sciatic nerve injury in *Sarm1-/-* mice, blood-borne Mɸ rapidly accumulate at the injury site but fail to accumulate in the distal stump^6^. Further, while monocytes will enter the distal stump, they fail to mature into Mɸ^5^. This suggests that a SARM1-dependent injury-induced Mɸ response could play a role in Wallerian degeneration after injury.

It remains unclear whether loss of *sarm1* in other non-neuronal cell types plays a role in the inflammatory cascade, or if the dampened inflammatory response in our peripheral nerve injury model^6^ is an indirect effect of *sarm1* loss in neurons. Mɸ respond to damage associated molecular patterns (DAMPs) that are released from the injury site and distal stump of the nerve in response to injury. Since the nerve fails to degenerate in *Sarm1-/-* mice, there is a reduction in the release of DAMPs. which could be too low to trigger a full inflammatory response. However, it is also possible that SARM1 activation in Mɸ themselves is required to activate the appropriate degeneration induced inflammatory cascade, including immune cell recruitment. Work from the Bowie group has recently reported that SARM1 is required for oxidative phosphorylation and glucose metabolism in cultured Mɸ^10,11^, indicating that SARM1 is required for Mɸ homeostasis. Additionally, data from our group has shown that *Sarm1-/-* delays regeneration, which could be due to changes in Mɸ activation in response to injury. In order to parse out the role of SARM1 in activated Mɸ states, we performed a series of physiological assays in wildtype and global *Sarm1-/-* splenic Mɸ. We found that loss of *sarm1* impairs Mɸ polarization and transition between inflammatory states *in vitro*. Our data suggest that SARM1 is necessary for the inhibition of “M2” signaling pathways, meaning that *Sarm1 -/-* Mɸ are in a constant state of “M2” activation, thus preventing them from fully transitioning to an “M1” phenotype. Importantly, we found that *Sarm1-/-* Mɸ are unable to support regeneration in a peripheral nerve injury model. Mɸ specific *Sarm1* conditional knockout mice further support that Mɸ require SARM1 to respond to peripheral nerve damage, including clearing myelin debris. Interestingly, neuronal specific *Sarm1* conditional knockout mice do not phenocopy germline *Sarm1-/-* mice, and surprisingly have similar myelin clearance to WT. Together, our data suggests that SARM1 is necessary for Mɸ functionality and is essential in driving the Mɸ response to axonal damage.

## Materials and Methods

### Experimental Design and Statistical Analyses

#### Animals

All procedures involving mice were approved by Ball State University and The University of South Carolina Animal Care and Use Committee (IACUC) and were performed in accordance with guidelines developed by the National Institutes of Health. Adult (8-23 week-old) male and female mice on a C57BL/6 background were used throughout the study. Mice were housed under a 12-hour light/dark cycle with standard chow and water ad libitum. Mouse lines included: C57BL/6 (Jackson Laboratories, Stock No. 000664), *Sarm1-/-* (Jackson Laboratories, Stock No. 018069), *ROSA26-tdTom* constitutively expressing membrane bound tdTomato in all cells (Jackson Laboratories, Stock No. 007576), and an in-house generated *ROSA26-tdTom;Sarm1-/-* line to constitutively express membrane tdTomato in *Sarm1-/-* cells. Conditional knockout lines were generated by crossing *Sarm1 fx/fx* mice (a kind gift from Dr. Jing Yang at Peking University) with B6.129P2-*Lyz2^tm1(cre)Ifo^*/J: (Jackson Laboratories, Stock No 004781) for specific knockout of *sarm1* in macrophages or with B6.Cg-Tg(Syn1-cre)671Jxm/J: (Jackson Laboratories, Stock No 003966) for specific knockout of *sarm1* in neurons.

#### Genotyping

Genomic DNA was obtained by ear biopsies and boiled at 100°C for 20 min in 100µl of alkaline lysis buffer (25 mM NaOH and 0.2 mM EDTA in ddH_2_0) and neutralized with 100 µl of 40 mM Tris-HCl (pH5.5). For PCR genotyping 5µl 5X Bio Rad GoTaq Buffer (Promega #M791A), 2.5µl MgCl2 buffer (Promega #M791A), 2µl of primer mix, 2µl of DNA sample, 0.5µl of dNTPs (Promega #U151B), and 0.2µl of GoTaq enzyme (Promega #M300) and ddH_2_0 was added to a total volume of 25 µl per sample. Cycling conditions and product sizes are listed in the table below:

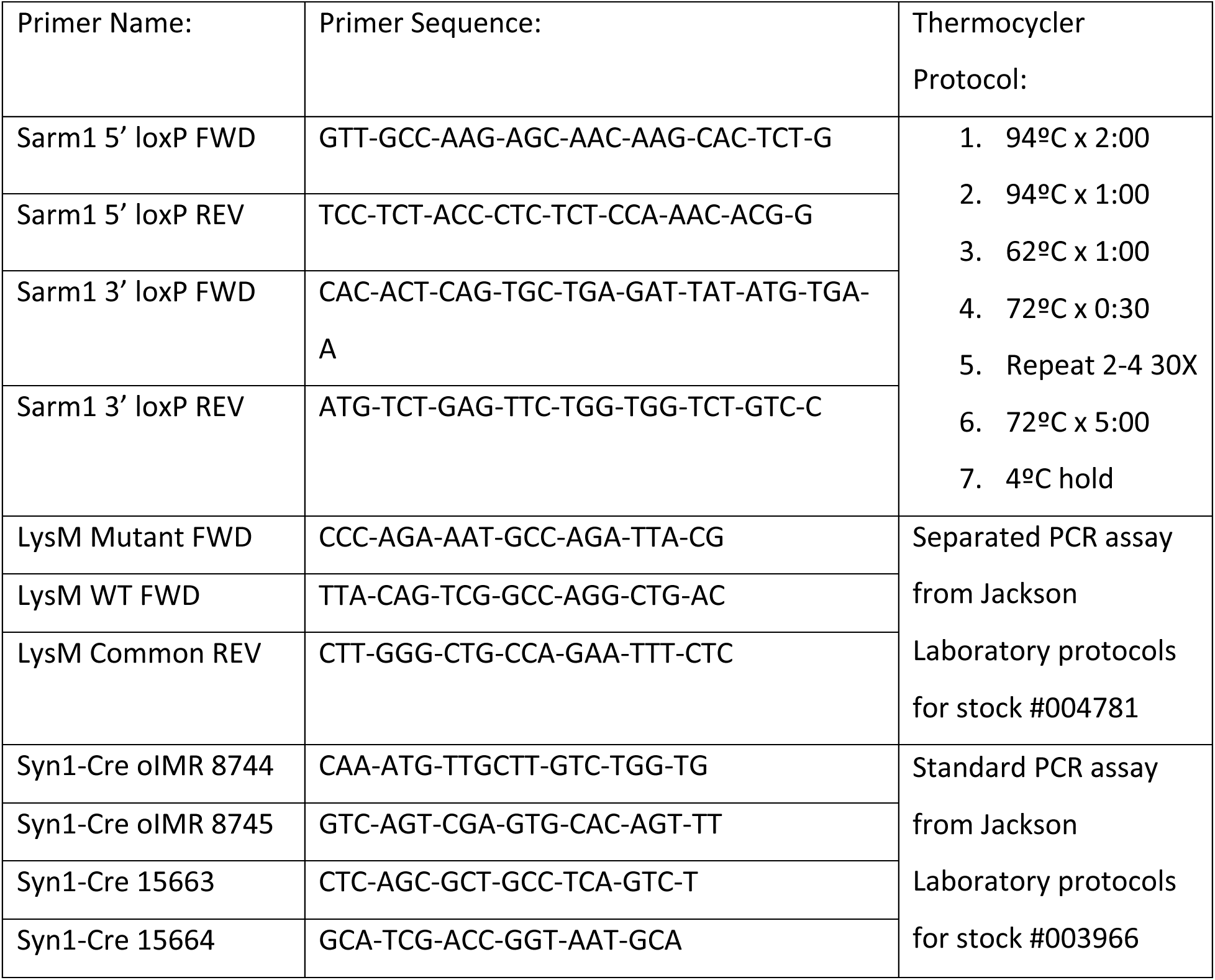

#### Sciatic Nerve Injections

Adult, male and female, (12 weeks old) wildtype mice were used for sciatic nerve injections. Surgeries were carried out under aseptic conditions. Mice were deeply anesthetized using 5% isoflurane and maintained with 3-3.5% isoflurane (SomnoSuite, Kent Scientific). Buprenorphine (0.1mg/kg or 3.25mg/kg of Extended-Release Buprenorphine) was given pre- and post-operatively. Thighs were shaved and disinfected with 70% ethanol (Fisher Scientific #22-363-750) and iodine (Med Vet International PVP-PAD). An incision was made at mid- thigh and underlying muscles were separated, exposing the sciatic nerve. The incision was held open using a retractor (FST #17005-04). Curved Dumont forceps (FST #11271-30) were gently inserted underneath the nerve to induce tension and provide a downward angle for injection. The forceps were held in place with a mosquito hemostat (FST #13008-12). For sciatic nerve crush, the nerve was crushed for 15 seconds, using fine forceps (Dumont #55, Roboz Surgical Instruments, #RS-5063). 1.0 µL of Mɸ cell suspension (∼50,000 cells) was slowly injected into the nerve, distal to the injury site using a 10 µl NanoFil syringe with a 36-gauge beveled needle (World Precision Instruments #NF36BV-2). Mɸ were prepared from spleens from *ROSA26-tdTom* or *ROSA26-tdTom;Sarm1-/-* mice as described below and resuspended in sterile PBS for injection.

#### Behavior

Mice aged 10-20 weeks at time of surgery were used for behavioral analysis. Mice used for behavioral analysis were moved to the testing room 24 hours before initiating testing. Post-surgical behavior and gait analysis were performed using a BlackBox Bio system with PalmReader software. Water was taken away 10 minutes before entering the BlackBox. Mice were then given 10 minutes to acclimate to the BlackBox. Mice were then tested for 20 minutes. Walking distance, rearing time, standing weight bearing, and toe spread ratio were analyzed after testing.

#### Cell Culture

All primary cell cultures were derived from *Sarm1-/-* or wild type male and female mice between 14-23 weeks of age.

##### Primary Mɸ Cultures

Spleens were harvested and passed through a 70 μm cell strainer. Cells were rinsed with Leibovitz’s medium (L-15) and centrifuged at 650x*g* for 5 minutes. Red blood cells were removed with ACK lysing buffer (Gibco #A10492-01). Cells were washed in (DMEM F-12 + Glutamax + 10% Fetal Bovine Serum (FBS)) and spun down at 650xg for 5 minutes. Cells were resuspended in complete Mɸ media (DMEM-10% FBS-0.05μg/mL mCSF) and plated onto 35mm plastic dishes. The cells were cultured at 37°C and 5% CO_2_ for 5 days with complete media change every other day.

##### Mɸ Polarization

On DIV 5, the culture media was replaced with treated media which included one of the following immunological stimuli: m-CSF (Abclonal #RP01216, 0.5µg/mL), IL-4 (Abclonal #RP01161, 0.1µg/mL), IFNɣ (Abclonal #RP01070, 0.1µg/mL), or lipopolysaccharide (Sigma #L6529, 0.1µg/mL), or a combination of both IFNɣ + LPS. The Mɸ were allowed to polarize for 24 hours before being re-plated over neurons (co-culture) or on PDL-treated coverslips (Mɸ only studies). For replating, adherent Mɸ were removed with 0.5% Trypsin. Cells were washed (DMEM F-12 + Glutamax-10% FBS-1x pen/strep) and centrifuged at 650xg for 5 minutes twice. For co-cultures, Mɸ were resuspended in neuronal media (DMEM F-12 + Glutamax-10% FBS-1x N1 supplement) For Mɸ only studies, Mɸ were resuspended in Mɸ complete media. For Mɸ injection surgeries, cells were suspended in sterile 1X PBS for a concentration of 5.0 x 10^7^cells/mL.

##### Neuronal Culture

Dorsal root ganglion (DRG) neurons were cultured from 8-12 week old *Sarm1-/-* or wildtype mice as previously described ^2^. Briefly, DRGs were digested in collagenase and dispase (1mg/mL each) for 25 minutes. Cells were triturated with fire-polished pipettes and washed in DMEM F-12 + Glutamax and 10% FBS and 1x Penicillin/Streptomycin (Gibco #15140-122) and centrifuged at 160xg for 5 minutes. Cells were resuspended in complete media (DMEM F-12 + Glutamax-10% FBS-1x N1 supplement) and plated onto glass coverslips. Coverslips were etched with 1M HCl before being coated in poly-l-lysine and 2µg/mL of laminin (Sigma #3400-010-01). Neurons were allowed to adhere for 24 hours before Mɸ were added. Co-cultures were fixed after 24 hours.

#### Immunostaining

##### Cell culture

Mɸ were stained on the sixth day of culture after having 24 hours to fully adhere to the coverslips after being re-plated. They were fixed with 4% paraformaldehyde at room temperature for 15-20 minutes, followed by two 5-minute washes in 1x PBS. The cells were then permeabilized with 0.3% Triton-X-100 in PBS for 10-15 minutes, which was followed by a 45-minute blocking step with cell blocking buffer (0.1% Triton-X-100/PBS-2%BSA-2%FBS). Primary antibodies were as follows: for Mɸ only cultures: rat α-F4/80 (Invitrogen #MA1-91124, 1:1000) and rabbit α-CD68 (Cell Signaling #97778S, 1:1000). For Mɸ-DRG co-cultures: rabbit α-CD68, mouse α-acetylated tubulin (Millipore #MABT868, 1:1000), and chicken α-NFH (Aves #NFH, 1:1000). For DRG only or media only DRG cultures: mouse α-acetylated tubulin and chicken α-NFH. For Mɸ myelin phagocytosis assay: rat α-F4/80.

Primary antibodies were incubated at 4°C overnight. The next day, they were washed 3 x 5 minutes with 1x PBS. Secondary antibodies were as follows: for Mɸ only cultures: α-mouse Cy5 (Jackson #715-175-150, 1:200), α-rat 488 (Jackson #712-545-150, 1:200), and α-rabbit 594 (Invitrogen R37119, 2 drops/mL). For Mɸ-DRG co-cultures: α-rabbit brilliant violet (Jackson #711-675-152, 1:150), α-mouse Cy5, and α-chicken Cy3 (Jackson #703-165-155, 1:200). For DRG only or media only DRG cultures: α-mouse Cy5 and α-chicken Cy3. For Mɸ myelin phagocytosis assay: α-rat 488.

Secondary antibodies were incubated at room temperature for 45-60 minutes followed by two washes in 1x PBS. The DRG-Mɸ co-cultures, media only neurons, and the DRG only cultures were also stained with actin green (Invitrogen #R37110, 2 drops/mL) for 10-15 minutes followed with two washes of 1x PBS. Mɸ myelin phagocytosis coverslips were stained with OilRedO (ThermoFisher #A1298914) following secondary antibody incubation^12^.

Coverslips were rinsed with milliQ water and mounted using ProLong^TM^ Gold antifade reagent (Invitrogen #P36930) or DAPI Fluoromount (SouthernBiotech #0100-20). The slides were allowed to dry at room temperature for 24 hours and were then stored at 4°C until imaging.

##### Tissue Sections

L4-6 DRGs and sciatic nerves were harvested from naïve and 7d post SNC mice. Tissues were fixed in 4% paraformaldehyde/1X PBS for 1 hour. DRGs were stored in 30% sucrose/1X PBS at 4°C for at least 24 hours. Samples were flash frozen with dry ice in Tissue-Tek O.C.T. Compound (Ref #4583) and sectioned at 14µm at −20°C and mounted on Superfrost Plus Gold slides (Fisherbrand #1518848). Samples were left to dry overnight at RT and stored at −20°C. Samples were washed 3 x 5 minutes with 1X PBS at RT and permeabilized with 0.3% Triton x 1X PBS for ten minutes. Samples were then blocked for 45 minutes in 5% donkey serum/0.1% Triton/1x PBS. Primary antibodies diluted in blocking buffer were then added: chicken α-NFH (Aves Labs #NFH 1:100), chicken α-NFM (Aves Labs #NFM 1:100), chicken α-NFL (Aves Labs #NFL 1:100), rabbit α-CD68, (Cell Signaling Technology #E307V 1:500), rabbit α-CD206 (Cell Signaling Technology #E6T5J 1:500), α-SCG10 (Novus Biologicals #NBP1-49461 1:500), α-Myelin Basic Protein (Bio Legend #808401 1:500), α-ATP8A2 (Invitrogen #PA5 −65256 1:500) and/or rat α-F4/80 (Invitrogen #MA1-91124 1:500) and slides were incubated at 4°C overnight. Samples were washed 3 x 5 minutes with 1X PBS at RT. Secondary antibodies diluted in blocking buffer: α-rat Cy3 1:250 (Jackson ImmunoResearch #712-166-150), α-chicken 488 1:250 (Jackson ImmunoResearch #703-546-155), α-rabbit Cy5 1:250 (Jackson ImmunoResearch #711-175-152), α-goat Cy3 (Jackson ImmunoResearch #705-166-147 and/or α-rat 488 1:250 (Jackson ImmunoResearch #712-545-150) incubated at RT for 1 hour. Slides were then washed 3 x 5 minutes with 1X PBS and once in DI water. Slides were then left to dry for 5 minutes. Cover slips (Fisherbrand #12541023) were mounted with Fluoromount with DAPI (SouthernBiotech #0100-20) and left to dry at RT overnight. Slides were stored at 4°C until used.

#### Western Blotting

Cells and tissues were lysed with RIPA buffer (150 mM NaCl, 50 mM TrisHCl, 1% NP-40, 3.5 mM sodium dodecyl sulfate, 12 mM sodium deoxycholate, pH 8.0) supplemented with 50 mM β-glycerophosphate (Sigma-Aldrich, #G9422-100G), 1 mM Na3VO4 (Sigma-Aldrich, #S6508-10G), and 1:100 protease inhibitor cocktail. Lysates were centrifuged at 14,000 rpm at 4°C for 15 min, supernatants were collected and protein concentrations were measured with the Pierce BCA protein kit (Thermo Scientific, #23227) using a SpectraMax at 562nm (Molecular Devices SpectraMax i3). Samples were diluted with 2x Laemmli sample buffer (Bio-Rad, #1610737) containing 5% β-mercaptoethanol (EMD Millipore, #6010), boiled for 10 min at 98°C, and stored at −20°C. For SDS-PAGE, equal amounts of total protein (15-50µg) were loaded for separation in a 12% gel. Proteins were transferred onto Nitrocellulose membrane (EMD Millipore, #IPVH00010) for 1h at 100V or on PVDF membrane (‘BioRad #162-0177) for 2.5hr at 200mA in ice-cold transfer buffer (25 mM TrisHCl, 192 mM Glycine, 10% Methanol). Membranes were blocked in 5% BSA (BioRad, #1706404) prepared in 1X TBS-T (TBS pH 7.4, containing 0.1% Tween- 20) or blotting-grade blocker (Biorad, #1706404) for 1 h at room temperature, and probed overnight at 4°C with the following primary antibodies diluted in 1X TBS-T with 2-3% BSA (Fisher Scientific, #BP1600): α-CD68 (Cell Signaling Technologies, #97778), α-ERK1/2 (Cell Signaling Technologies, #4695), α-pNFkB (p65) (Cell Signaling Technologies, #3033), α-NFkB (p65) (Cell Signaling Technologies, #8242), α-pSTAT6 (Cell Signaling Technologies, #9361), α-STAT6 (Cell Signaling Technologies, #9362), α-Arginase-1(Cell Signaling Technologies, #93668), α-CD14 (Cell Signaling Technologies, #93882), α-CD206 (Cell Signaling Technologies, #24595), Dectin-1 (Cell Signaling Technologies, #30260), α-GAPDH (Cell Signaling Technologies, #5174), α-iNOS (Invitrogen, #PAI-038), α-SARM1 (BioLegend, #696602, 1:250). The following Horseradish peroxide (HRP)-conjugated secondary IgG antibodies were used from Cell Signaling Technologies: α-rabbit (7074), α-rat (#7076P2), α-mouse (#7076). Membranes were incubated in (HRP)-conjugated secondary IgG antibodies diluted in 1X TBS-T with 2-3% BSA for 1h at room temperature. The HRP signal was developed with chemiluminescent substrates (Thermo-Fisher Scientific West Pico, #34577 or Femto, #34095; Millipore #WBKLS0100).

#### Quantitative PCR

Cultured Mɸ were polarized on DIV5 and lysed on DIV6 for RNA (Qiagen RNeasy Micro Kit #74004). Complementary DNA was reverse transcribed (BioRad iScript kit #1708891) and was prepared in a 96-well hard-shell PCR plate (BioRad, #HSP9655) at a concentration of 5ng with SYBR Green Supermix (BioRad, #1725271) at 1X concentration, forward and reverse primers at 500nM **(Table S1)**, and nuclease-free water. The plate was run on a BioRad CFX Maestro thermocycler at the recommended settings for the Supermix. Data was normalized to 12S and analyzed for fold change relative to the control condition mCSF.

#### Mɸ Assays

##### Scratch Assays

Polarized Mɸ were seeded at ∼40,000 to 50000 cells/well in an Imagelock 96-well Plate (Sartorius #BA-04855) and incubated at 37°C for 18h-20h. The scratch was made using the Incucyte 96-well Woundmaker Tool. Each well was washed twice with complete media (DMEM, 10% FBS, and m-CSF). Fresh complete media were added after washing, and the plates were placed into the Incucyte Live-Cell Analysis System. The cells were scanned/imaged every 2h in the live-cell analysis software (Scratch Wound and Wide mode settings, 10X objective).

##### Invasion Assays

Imagelock 96-well Plate (Sartorius #BA-04855) was coated with a thin layer of Matrigel Matrix (100ng/ml) (Corning #354277). Polarized Mɸ were seeded at ∼40,000 to 50000 cells/well in the Imagelock 96-well Plate and incubated at 37°C for 18h-20h. A scratch was made using the Incucyte 96-well Woundmaker Tool. Each well was washed twice with complete media (DMEM, 10% FBS, and m-CSF). Fresh complete media were added after washing, and the plates were cooled at 4°C for 5 minutes. The cells were overlayed with a top layer of Matrigel Matrix (4mg/ml) and incubated at 37°C for 30 minutes. Additional media with or without recombinant mouse complement component C5a (R&D system, #2150-C5/CF) was added after incubation. The plates were placed into the Incucyte Live-Cell Analysis System, and the cells were scanned/imaged every 2h in the live-cell analysis software (Scratch Wound and Wide mode settings, 10X objective).

##### NAD/NADH- Glo Assay

Cultured Mɸ were seeded at ∼15,000cells/well in a 384-well white flat-bottom microplate and incubated at 37°C for 1h-24h. The NAD/NADH-Glo detection reagent (Promega, #G9072) was prepared according to the manufacturer’s instructions. An equal volume of reagent was added to each sample/well and incubated at room temperature for 30 minutes. The luminescence was measured using SpectraMax (Molecular Devices SpectraMax i3).

##### Proliferation and Apoptosis

Mɸ were cultured in a 96-well plate at a density of ∼150,000 cells/ well. Incucyte Nuclight Rapid red reagent (Essen Biosciences, #4717) and Incucyte caspase 3/7 green dye (Sartorius, #4440) were added to each well at a 1:1000 dilution, respectively. Cells were scanned/ imaged at 2h interval in the live-cell analysis software.

##### Repolarization Assay

Cultured Mɸ were seeded at ∼10mil cells/well in a 6-well clear-bottom cell culture plate and were polarized on DIV5 with either IL-4 or a mixture of IFNɣ and LPS. On DIV6, some cells were re-plated into a 12-well clear-bottom cell culture plate at ∼1mil cells/well. On DIV7, the cells were either polarized with the opposite immunological stimuli or given plain media. On DIV9, the cells were lysed for proteins.

##### Myelin Phagocytosis Assay

Cultured Mɸ were polarized with mCSF, IL-4, or LPS on DIV5 and seeded at ∼500,000cells/well on PDL-coated glass coverslips in a 24-well clear-bottom culture plate on DIV6 as described for Mɸ culture. The cells were given a 1% w/v solution of peripheral nerve myelin as prepared in^12^ on DIV7. For no clearance, the cells were fixed and stained on DIV8. For clearance, the cells were washed with complete media on DIV8 and fixed DIV9.

#### Imaging

Images were acquired on a Zeiss AxioObserver 7 microscope with an AxioCam 807 monochrome camera with a 20x (Plan Apochromat 20x/0.8 air) objective (for macrophage cultures) or a 40x (Plan Apochromat 40x/1.3 Oil) objective (for tissue sections). DRG tissues were imaged using the Apotome 3 with a z-step size of 0.5 µm. An Excelitas X-cite mini+ lamp was used for fluorescent channels and adjusted by channel. Lamp intensity was set to 50% for tissue sections and 60% for cell culture samples. For neurite length images a Zeiss AxioObserver 7 microscope with an AxioCam 503 monochrome camera with a 10x (PH1 10x/0.3 air) objective was used.

##### Imaging

Images used for full-nerve Scg10 and F4/80 analysis were taken on a Zeiss AxioObserver 7 microscope with an AxioCam 807 monochrome camera. Images were acquired with a Plan-Apochromat 20x/0.8 M27 air objective. Conventional fluorescence was used and z-projections were formed with a 2µm step size. For quantification, immunofluorescence was measured every 500µm beginning 500µm proximal to the injury site and ending 1mm distal to the injury site. Scg10 exposure was set to 0.72s and F4/80 was set to 0.23s. Lamp intensity was set to 60% for all channels.

Images taken for MBP/F4-80/ATP8A2 high resolution quantification were taken on a Zeiss AxioObserver 7. A Zeiss 40x/1.3 Oil DIC (UV) VIS-IR 420762-9800 objective was used to take images at the injury site, 1mm distal to the injury site, and the proximal stump of a naïve wild-type nerve. Conventional fluorescence was also used, and z-projections were formed with a 0.8µm step size. MBP exposure was set to 0.082s, F4/80 exposure was set to 0.036s, and ATP8A2 exposure was set to 1.8s. Representative images were taken on a ZEISS LSM 990 confocal microscope with a Plan-Apochromat 63x/1.4 NA Oil objective (420782-9900-799). AF647 (Cy5) was set to 4% power, Cy3 was set to 1% power, and AF488 was set to 0.5% power. Z step size was 0.5µm. 3D Projections that underwent LSM plus deconvolution were used for representative images.

#### Quantification

The longest neurite from at least 100 individual neurons in each condition was quantified using ImageJ or Neuromath (WIS) ^13^ by tracing from the cell body to the tip of the neurite. Neurons were identified by neurofilament heavy immunoreactivity. CD68 expression was measured in F4/80+ cells using ImageJ. At least 400 Mɸ were quantified from each condition. To quantify Mɸ phenotypes, cells were segregated based on size and shape from phase-contrast images. Only those cells that were also positive for F4/80 immunoreactivity were included in analysis. Phenotype parameters were as follows: small Mɸ were round with a diameter less than 11 µm. Large Mɸ had to have a diameter greater than 40 µm and a roughly circular cytoplasmic projection. Mɸ were considered elongated if they had a comet shaped projection in 1 or 2 directions making the Mɸ longer than wider. Mɸ were considered stellate if they had 3 or more distinct cytoplasmic projections. For each condition, at least 400 Mɸ were quantified across four independent cultures. The acetylated tubulin was quantified in ImageJ-Fiji by outlining and measuring mean pixel value 50 µm directly proximal to the cell body, the growth cone, and then 50 µm directly proximal to the growth cone (distal stump). The myelin phagocytosis assay was quantified in ImageJ-Fiji using mean pixel value relative to the area of each cell.

#### Statistical Analysis

Outliers were removed from data sets using the ROUT and Q=1% method from Prism (Graphpad). qPCR data and neurite length analyses were assessed for normality and then analyzed with a one-way ANOVA and multiple comparisons with Tukey post-hoc analysis, or a Kruskal-Wallis test. For qPCR heatmaps, data was log transformed (Y=log(Y)) and then analyzed by a two-way ANOVA with a Fishers LSD test.

Invasion and scratch assays, cell counts, confluency, caspase, proliferation, and behavioral data were analyzed by a two-way ANOVA with repeated measures. Sphericity was not assumed therefore a Geisser-Greenhouse correction was used. For a full model analysis, a Fishers LSD or Dunnett’s post-hoc analysis was used for multiple comparisons. For a main effects model analysis only, a Tukey-post hoc analysis was used for multiple comparisons.

Immunological stimuli polarized and repolarized western blots were analyzed by a two-way ANOVA. A full model analysis was performed with a Fishers LSD test.

NAD/NADH glo assays were analyzed by a two-way ANOVA. A full model analysis was performed with a Tukey post-hoc analysis for multiple comparisons.

Mɸ morphology was analyzed by a Chi-Square analysis comparing either effect of genotype on morphology or immunological stimuli treatment on morphology (independent of genotype).

## Results

### Loss of sarm1 reduces Mɸ accumulation following injury

Macrophages (Mɸ) are resident to both the sciatic nerve and dorsal root ganglia (DRG) but are present in very low numbers **(Fig1A,D,F)**. We previously showed that germline deletion of *sarm1* (*Sarm1-/-*) does not impair Mɸ or immune cell composition in adult sciatic nerves^6^ (**Fig1A,B**). Western blot analysis of sciatic nerves after injury (SNC) shows no significant decrease in myeloid cells or Mɸ by CD11b and CD68 expression, respectively (**Fig1A-B**). However, consistent with our previous work, while the injury site shows similar total macrophage accumulation, there is a decrease in the distal stump, as seen by F4/80, a Mɸ-specific surface marker^5,6^ (**Fig1C**). Further, we see a decrease in CD68+ Mɸ in the distal stump of *Sarm1-/-*mice, which could indicate a reduction in “activated” Mɸ (**Fig1C**). Work from the Giger lab has shown that Mɸ also respond in the lumbar DRGs following SNC by altering their cellular morphology, instead of an recruiting additional infiltrating myeloid cells^2^. To determine if loss of *sarm1* impacts Mɸ responses at the DRG, we lysed lumbar level DRGs before and after injury and probed for myeloid cells and Mɸ with CD11b and CD68, respectively (**Fig1D,E**). Interestingly, we did not see any difference in WT and *Sarm1-/-* animals. We then analyzed tissue sections to examine Mɸ morphology and activation state (**Fig1F**). We found that in naïve DRGs, Mɸ look similar morphologically in both WT and *Sarm1-/-*. However, 7d after SNC we see a qualitative reduction in expansion of Mɸ volume by F4/80 staining in the *Sarm1-/-* animals (**Fig1F**). Further, *Sarm1-/-* display an increase in CD163+ positive Mɸ in DRGs 7d post SNC compared to WT (**Fig1F**), suggesting that *Sarm1-/-* Mɸ are still in a homeostatic state. Together with our previous work^6^, the reduction in Mɸ accumulation and/or activation in the *Sarm1-/-* mice suggests an important role for SARM1 in Mɸ activation. To parse this out, we turned to primary Mɸ cell cultures.

**Figure 1:**
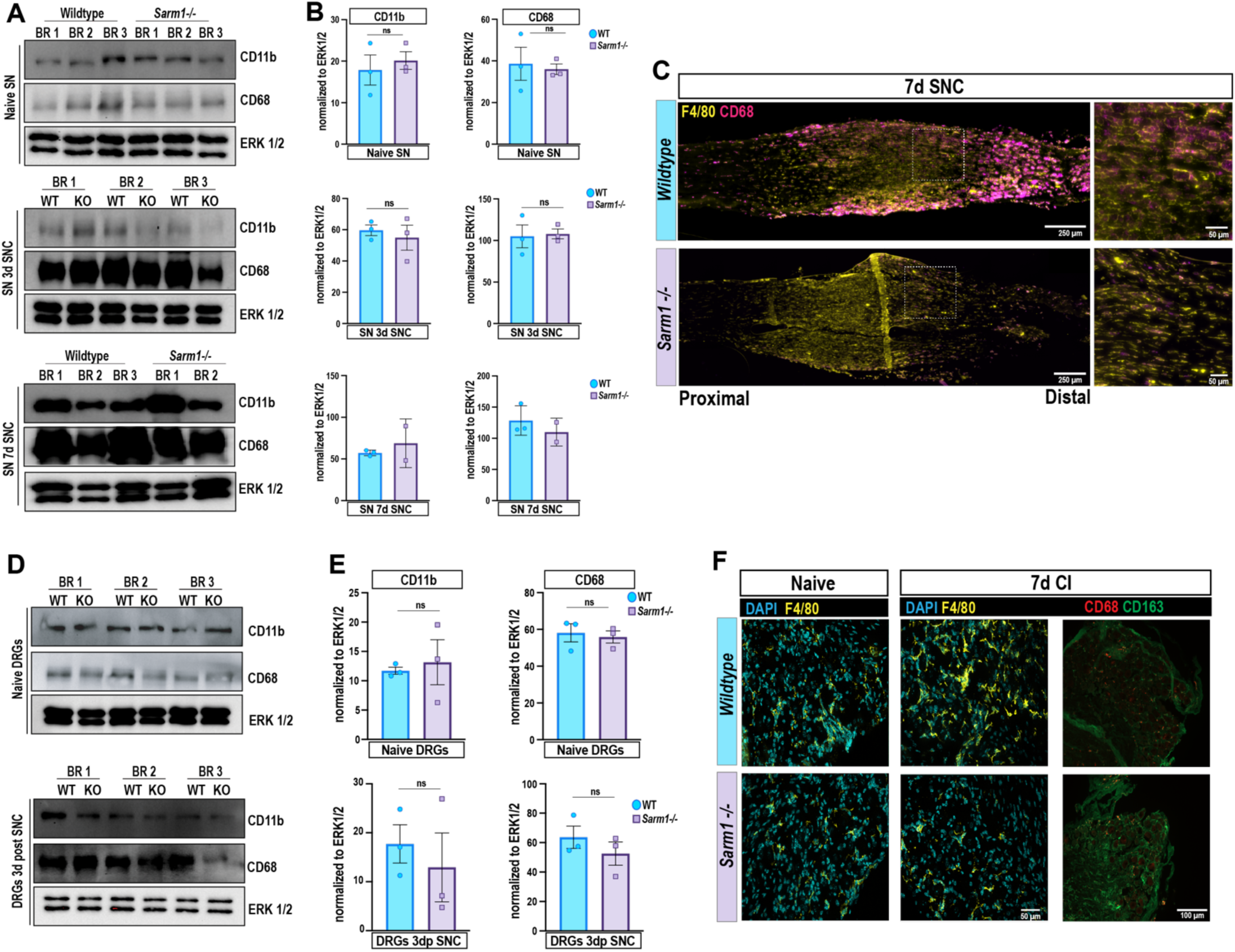
Mɸ loss is restricted to distal nerves in *Sarm1-/-* mice following peripheral nerve injury. (**A**) Western blots of whole sciatic nerves (SN) from Naïve, 3d, and 7d post sciatic nerve crush (SNC). Blots were probed for CD11b and CD68. ERK1/2 was used as a loading control. (**B**) Quantification of CD11b (left) and CD68 (right) from western blots in **A**. Expression was normalized to ERK1/2. Error bars = SEM for Naïve and 3d and Stdev for 7d. Each data point represents a biological replicate. ns = Not significant by nonparametric two-tailed student t-test. (**C**) Representative image of sciatic nerves 7d post sciatic nerve crush in WT (top) and *Sarm1-/-* (bottom). Tissues were stained for Mɸ markers (F4/80- yellow; CD68- magenta). Boxes are indicating the enlarged images. N= 3 biological replicates. Scale bar = 250 µm or 50 µm. (**D**) Western blots of lumbar dorsal root ganglia (DRG) from Naïve and 3d post sciatic nerve crush. Blots were probed for CD11b and CD68. ERK1/2 was used as a loading control. (**E**) Quantification of CD11b (left) and CD68 (right) from western blots in **D**. Expression was normalized to ERK1/2. Error bars = SEM. Each data point represents a biological replicate. ns = Not significant by nonparametric two-tailed student t-test. (**F**) Representative images of naïve or 7d post sciatic nerve crush (CI) DRG sections stained for Mɸ markers (F4/80-yellow; CD68-red; CD163-green) and nuclei (DAPI-blue) in WT (top) and *Sarm1-/-* (bottom). N= 3 biological replicates. Scale bars = 50 µm or 100 µm.

### SARM1 is expressed in mouse primary Mɸ

We chose to polarize Mɸ from the spleen due to their abundance and sensitivity to immunological stimuli *in vitro*. To determine the optimal time to treat the Mɸ, we assessed morphology at different time points after culturing (**Figure S1A**). Initially after plating, most Mɸ were still non-adherent and in suspension. These cells were not included in the total cell counts, but were included in the confluency analysis. By 5 days, however, most cells were adherent and > 50% confluency (**Figure S1C**). Thus, we determined that 5 days *in vitro* (DIV) was an optimal time to treat the Mɸ.

We cultured Mɸ for 5 days in m-CSF and 10% FBS to ensure we reached a homeostatic environment. On day 5, we supplemented media with either IL-4 to polarize these Mɸ towards anti-inflammatory, or IFNɣ or lipopolysaccharide (LPS), to polarize towards pro-inflammatory **(Fig2A)**. m-CSF + 10% FBS was used as a control medium. We polarized Mɸ for 24 hours with immunological stimuli and induced stress by replating. Purity of cultures was confirmed by F4/80 or CD68 **(Fig2C-E)**. Mɸ were then analyzed 24 hours after replating **(Fig2A)**. SARM1 was readily detectable in Mɸ regardless of prior immunological stimulation **(Fig2B)**. This is consistent with recent reports in cultured bone-marrow derived Mɸ treated with LPS or IL-4^10,11^. We further confirmed SARM1 expression at the protein level by western blot and at the transcript level by qRT-PCR **(Fig2D-G)**. Notably, there was a trending increase in expression at the protein level in the LPS conditions, but variability at the RNA level in both IL-4 and LPS conditions **(Fig2F,G)**. These data support recent reports that SARM1 is expressed in cultured Mɸ.

**Figure 2:**
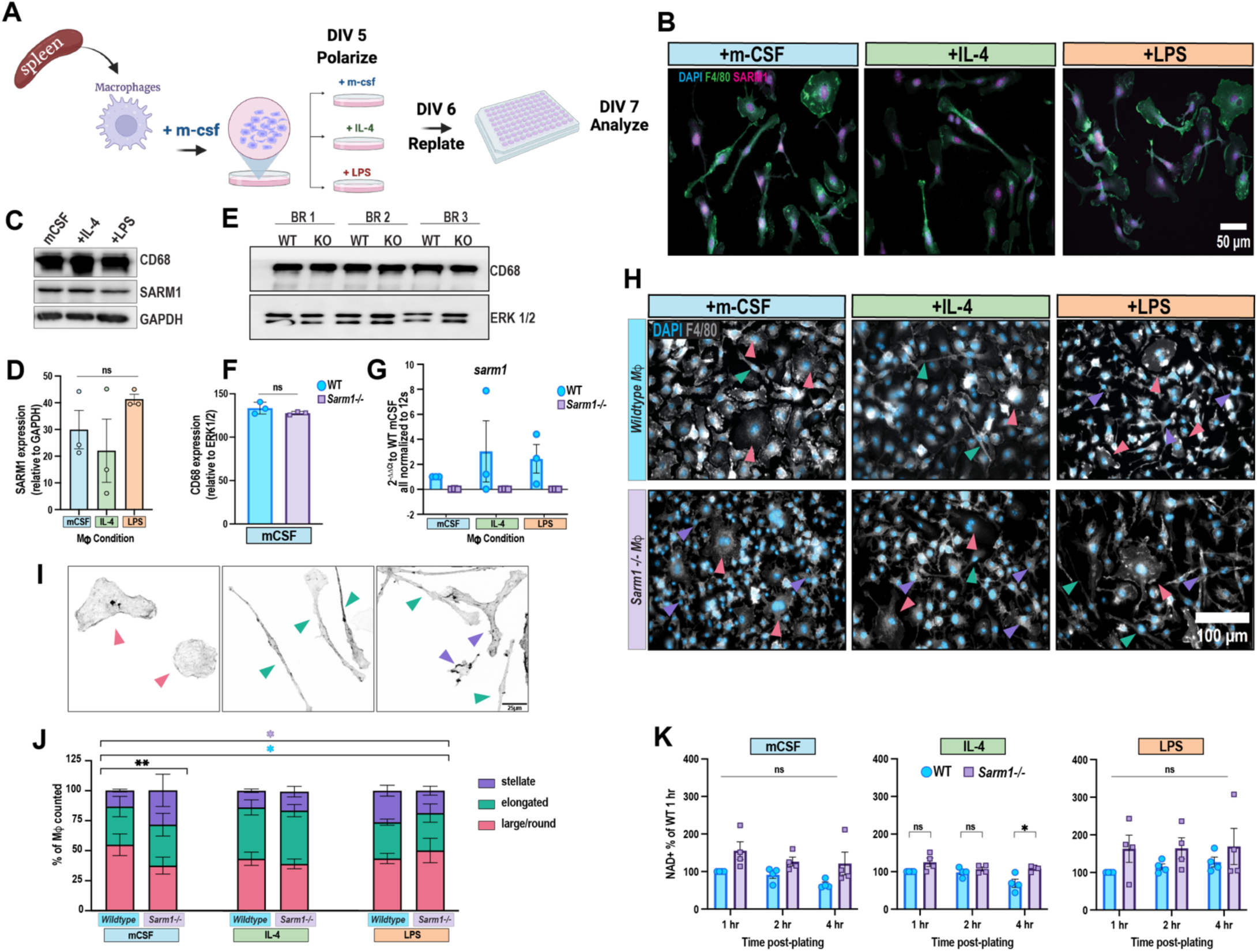
Mɸ express SARM1 and respond to immunological stimuli. (**A**) Diagram depicting primary splenic Mɸ culture and immunological stimulation to drive immunophenotypes. Made in BioRender. (**B**) Representative images of wildtype Mɸ cultured for 7 days and stained for F4/80 (green) and SARM1 (magenta). Cell nuclei stained for DAPI (blue). Scale bar = 50µm. (**C**) Representative western blot of cytokine stimulated wildtype Mɸ probed for CD68 and SARM1. GAPDH was used as a loading control. **D**) Quantification of SARM1 expression from (**C**) normalized to GAPDH. N = 3 biological replicates. Error bars = SEM. ns = not significant by Kruskal-Wallis with Dunn’s correction for multiple comparisons. (**E**) Representative western blot for CD68 in wildtype (WT) and *Sarm1-/-* (KO) Mɸ cultures. ERK1/2 was used as a loading control. N = 3 biological replicates (**F**) Quantification of CD68 expression from (**E**) normalized to ERK1/2. N = 3 biological replicates. Error bars = SEM. ns= not significant by nonparametric student t-test. (**G**) qRT-PCR of *sarm1* expression in stimulated WT and *Sarm1-/-* Mɸ. 12s was used as an internal control. Graph represents 2^ΔΔCt as compared to WT mCSF. Error bars = SEM. (**H**) Representative images of WT and *Sarm1-/-* Mɸ stained for F4/80 (grey) and DAPI (blue). Colored arrows indicated different Mɸ morphology as it corresponds to **I,J**. Scale bar = 100µm. (**I**) Example images of Mɸ morphologies; pink indicates large/round, green indicates elongated, and purple indicates stellate. Scale bar = 25 µm. (**J**) Quantification of Mɸ morphologies from Mɸ in **H**. N = 5 biological replicates, and ≥100 Mɸ per biological replicate were included in the analysis. Error bars = SEM. *p<0.05;***p<0.01 per Chi-squared analysis. (**K**) Quantification of NAD+/NADH Glo assay in stimulated WT and *Sarm1-/-* Mɸ. Data is represented as percentage of WT 1-hour levels. N = 3 biological replicates. Error bars = SEM. ns= not significant;*p<0.05 per two-way ANOVA with Tukey post-hoc test.

### Loss of sarm1 alters Mɸ response to microenvironmental signaling

We found that our culture paradigm led to distinct morphological differences in the Mɸ populations, indicating that polarization by immunological stimuli led to changes in Mɸ immunophenotypes (**Fig2H-J**). Assessment of F4/80+ Mɸ by phase contrast showed 3 distinct Mɸ phenotypes: Large/round, elongated, and stellate (**Fig2I**). Small Mɸ were considered to be uniformly round monocytes that have not matured into Mɸ and were therefore excluded from downstream morphological analysis (**FigS1A,B)**. Large and round Mɸ were classified by extensive cytoplasm and a defined F-actin cytoskeleton. These are typically phagocytic and considered to be pro-inflammatory. Elongated Mɸ were classified by F-actin protrusions extending unilaterally or bilaterally. Elongated Mɸ are typical of pro-regenerative or wound healing Mɸ^14^. Stellate Mɸ were defined as ratified or star shaped (**Fig 2I**). As previous work indicates that stellate Mɸ may support neuronal growth or prevent cell death following SNC^2^, so we considered them to be an anti-inflammatory phenotype. Chi-squared analysis of cell morphology phenotypes showed significant differences across immunological stimuli for both WT and *Sarm1-/-* (**Fig2J**). Consistent with previous reports, IL-4 treatment resulted in the most defined elongated Mɸ phenotype (**Fig2H,J**). LPS treatment led to an increase in stellate Mɸ compared to both mCSF and IL-4 treatment. Interestingly, at baseline conditions (treatment with mCSF), we found the most significant difference between WT and *Sarm1-/-* Mɸ morphologies (**Fig2J**). Consistent with the increased CD163 expression of *Sarm1-/-* Mɸ in DRGs after SNC in **Fig1**, *Sarm1-/-* Mɸ showed increases in stellate and elongated morphologies regardless of immunological stimuli stimulation, suggesting that loss of *Sarm1* leads to more alternatively activated or anti-inflammatory Mɸ phenotypes **(Fig2H,J)**.

As previously stated, SARM1 is a unique toll-like adapter protein due to its potent enzymatic activity. When activated, SARM1 breaks down NAD+ leading to a production of NMN and cADPR^15^. NAD+ levels impact cellular metabolism, and other groups have shown that bone-marrow derived Mɸ cultured from *Sarm1-/-*mice have an increase in glycolysis and oxidative phosphorylation^10^. Therefore, it is possible that Mɸ metabolism either regulates immunophenotypic changes or is altered during polarization and activation.

To determine if altered morphologies of *Sarm1-/-* splenic Mɸ are due to an accumulation of NAD+ levels, we performed an NAD/NADH Glo assay and measured NAD/NADH levels at 1-, 2-, and 4-hours following replating of 24 hour stimulated Mɸ **( (Fig2K)**. While we found a reduction in NAD+ consumption in all stimulation conditions in the *Sarm1-/-* Mɸ, there was little consumption of NAD+ in the WT cells at these time points **(Fig2K)**. Notably, there was only a significant difference in consumption between genotypes 4 hours after replating in the IL-4 conditions **(Fig2K** These data suggest that SARM1 dependent NAD+ metabolism is unlikely to alter Mɸ phenotypes in *Sarm1-/-* mice. To determine if phenotypic changes resulted from alterations in cell number, proliferation, or confluency we analyzed cultures over time and quantified total cell counts and confluency every 2 hours over the course of 6 days (**FigS1A-C)**. Cells were plated at a density of 1 million cells, however, only adherent cells are counted, so the total cell number is inaccurate for the first 24 hours (**FigS1B)**. While the total cell number increased rapidly between 24 and 48 hours for both WT and *Sarm1-/-* cells, there was a significantly higher WT cell count between 64 and 80 hours (**FigS1B)**. As Mɸ maturation continues through DIV 4 and 5, we see a continued trend of decreased cell counts in *Sarm1-/-,* but it does not reach statistical significance. Interestingly, while there is reduced total cell counts during this 5-day period for *Sarm1-/-* Mɸ, there is a significant increase in confluence between 68-88 hours, likely due to changes in overall cell morphology (**FigS1C).** However, by DIV 4 and 5 there is no difference between genotypes. Further, there is no difference in caspase mediated cell death indicated by nuclear expression of caspase 3/7 (**FigS1D,E**) or in cell proliferation indicated by a fluorescent nuclear dye (**FigS1F,G**). Together, this suggests that only Mɸ maturation and polarization are impacted by loss of *sarm1*. This is consistent with previous work showing delayed Mɸ maturation following sciatic nerve injury in *Sarm1-/-* animals^5^.

### Loss of Sarm1 alters Mɸ response to injury

The altered phenotypes of *Sarm1-/-* Mɸ suggest they might have altered functions. To determine if loss of *sarm1* impacts the functionality of Mɸ we subjected them to a series of injury assays that are relevant to physiological responses required following nerve injury. First, we subjected polarized Mɸ to a two-dimensional scratch wound assay (**Fig3A-C; S2A-D; Supplemental Movies 1-4**). Here, Mɸ were plated to near confluency (∼90%) and a scratch was induced using a WoundMaker tool that induces a uniform scratch of approximately 700-800 µm diameter was induced in all wells using a WoundMaker tool. Mɸ were then monitored by live imaging every 2 hours for a total of 48 hours (**Fig3A-C; S2A-D**). Regardless of genotype, IL-4 treated Mɸ showed the most wound closure as measured by relative wound density (**Fig3B,C; FigS2A-C**). A simple linear regression analysis to identify rate (slope) and density (elevation) showed significant differences between genotypes in elevation between genotypes for mCSF and IFNɣ conditions, and a significant difference in slope for LPS conditions, but no differences for IL-4 stimulation (**FigS2A-C**). These data suggest that immunological stimuli differentially impacts WT and *Sarm1-/-* Mɸ. Within group analyses further highlighted immunological stimuli differences (**Fig3B,C**). WT Mɸ display significant increase in relative wound density over time for mCSF, IL-4, and LPS conditions, but not IFNɣ (**Fig3B**). Additionally, IL-4 treated WT Mɸ showed significant increase in wound closure compared to mCSF conditions between 36-42 hours post scratch (**Fig3B**). *Sarm1-/-* Mɸ on the other hand displayed significant increases in relative wound density over time in only mCSF and IL-4 conditions (**Fig3C**). Further there was significant decrease in wound density in both LPS and IFNɣ conditions compared to mCSF between 6-34 hours and 4-30 hours, respectively, post scratch (**Fig3C**). Together, these data suggest that *Sarm1-/-* Mɸ may be more alternatively activated (anti-inflammatory) and primed for wound healing, and that IL-4 stimulation further enhances this phenotype. However, when *Sarm1-/-* Mɸ are classically activated (i.e. with LPS or IFNɣ), they fail to make a rapid transition during the scratch assay to fully close the wound.

**Figure 3:**
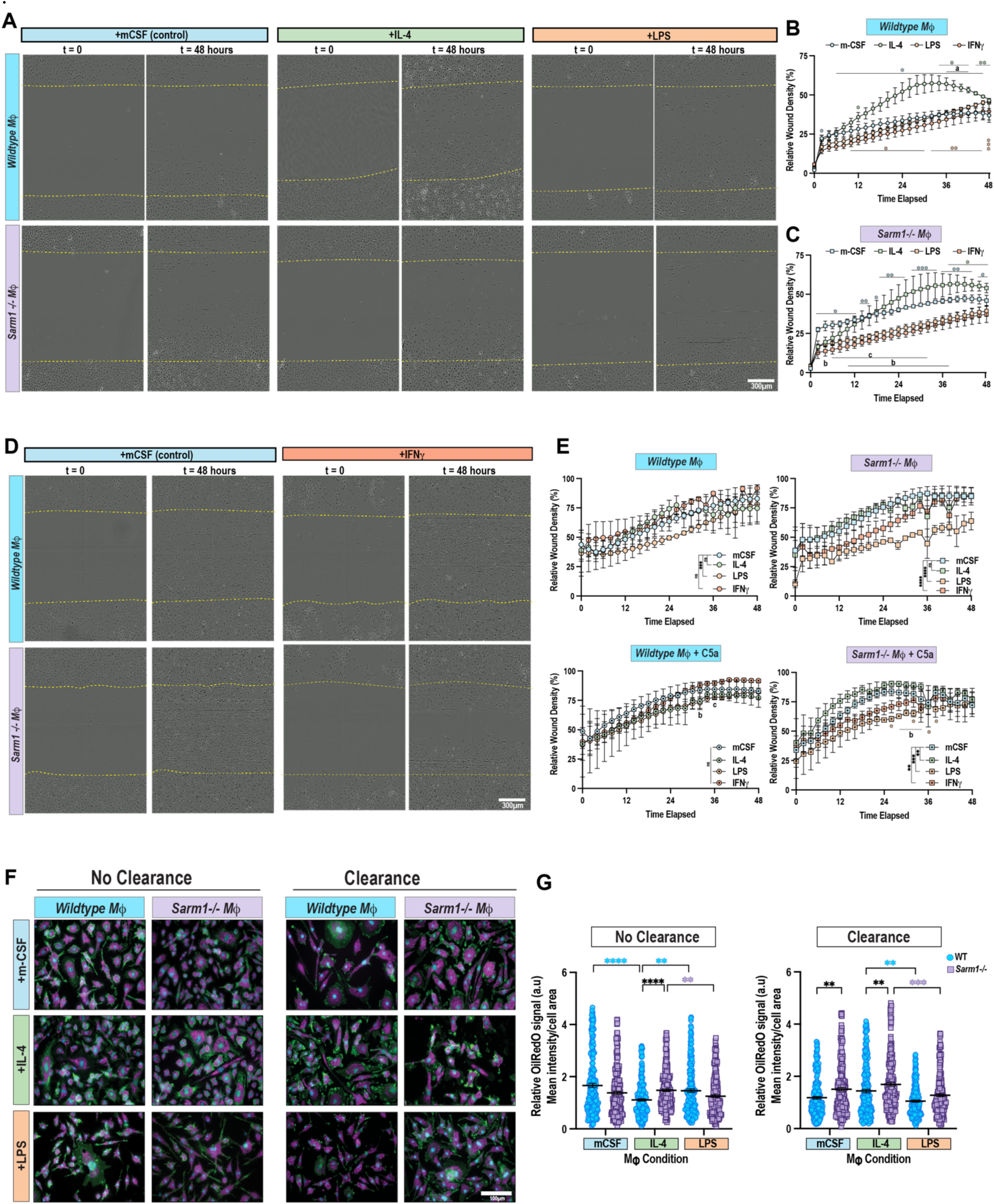
Loss of *sarm1* differentially impacts Mɸ response to stimuli. (**A**) Representative still images from live cell imaging sequences following a 2D scratch assay of WT and *Sarm1-/-* stimulated Mɸ. t=0 is immediately after the scratch and t=48 hours is 48 hours post-scratch. Scratch boundaries are indicated by yellow dashed lines. Scale bar = 300µm. (**B,C**) Quantification of the relative wound density of WT (**B**) or *Sarm1-/-* (**C**) Mɸ cultures after the scratch assay in **A** represented as a percentage of t=0. N=3 biological replicates. Error = SEM. *p<0.05; **p<0.01; ***p<0.005; letters (a,b,c) p<0.05. by two-way repeated measures ANOVA with Dunnett post-hoc test. Blue asterisk = mCSF compared to t=0. Green asterisk = IL-4 compared to t=0. Orange asterisk = LPS compared to t=0. a = mCSF compared to IL-4; b = mCSF compared to LPS; c = mCSF compared to IFNɣ conditions at indicated timepoints. (**D**) Representative still images from live cell imaging sequences following a 3D invasion assay in Matrigel of WT and *Sarm1-/-*stimulated Mɸ. t=0 immediately after the scratch and t=48 hours is 48 hours post-scratch. Scratch boundaries are indicated by yellow dashed lines. Scale bar = 300µm. (**E**) Quantification of the relative wound density of the invasion assay without C5a added (top) and with C5a added as a chemoattractant (bottom) represented as a percentage of t=0. N=2-4 biological replicates. Error = SEM. Letters (b,c) p<0.05; **p<0.01; ***p<0.005; ****p<0.001 by two-way ANOVA, main effects model with Tukey post-hoc test after simple linear regression model indicated slopes were not different. Wildtype Mɸ + C5a main effects could not be calculated because slopes were significantly different. Orange asterisk = LPS compared to t=0. b = mCSF compared to LPS; c = mCSF compared to IFNɣ conditions at indicated timepoints. (**F**) Representative images of F4/80 (green) positive Mɸ stained for Oil Red O lipids (magenta) following a myelin challenge assay. No clearance indicates a 24 hour myelin treatment and clearance indicates 24 hours of myelin treatment followed by myelin removal for 24 hours. Scale bar = 100µm. (**G**) Quantification of OilRedO intensity per Mɸ (normalized to Mɸ cell area). N = 3 biological replicates and each data point = 1 Mɸ. Mean +/- SEM is shown. Outliers were removed with ROUT Q=1. **p<0.01; ***p<0.005; ****p<0.001 by Kruskal-Wallis test with Dunn’s correction for multiple comparisons. Black asterisk = WT to *Sarm1-/-* comparisons. Purple asterisk = *Sarm1-/-* to *Sarm1-/-*comparisons. Blue asterisk = WT to WT comparisons.

We recently demonstrated that there is a decrease in Mɸ accumulation in distal nerves of *Sarm1-/-* mice after sciatic nerve crush^6^ (**Fig1B,D**). Since a majority of these Mɸ are derived from monocyte precursors from the blood^2^, we reasoned that *Sarm1-/-* Mɸ may have impaired invasion mechanisms. To test this, we seeded Mɸ on a layer of Matrigel before inducing a scratch wound (**Fig3D-E; FigS2 E-G; Supplemental movies 5-8**). We followed that by another layer of Matrigel so that the Mɸ were encased in a three-dimensional matrix during wound healing. Similar to the two dimensional scratch assay, *Sarm1-/-* Mɸ at baseline (mCSF condition) and treated with IL-4 performed better than WT Mɸ (**Fig3E**). Although, WT Mɸ also eventually showed full wound closure by 48 hours (**Fig3E**). However, when Mɸ were polarized with LPS or IFNɣ prior to the invasion assay, *Sarm1-/-* Mɸ significantly lagged behind WT, and failed to close the wound even at 48 hours (**Fig3D,E**). The addition of C5a, a potent chemoattractant, did rescue this deficit in the IFNɣ treated condition, but not in the LPS condition (**Fig3E; FigS2 F-G; Supplemental movies 9-12**). This further suggests that at baseline, *Sarm1-/-* Mɸ are more alternatively activated (“M2”), and that classical activation (“M1”) impairs their response to wound healing assays.

### Loss of Sarm1 impairs clearance of PNS myelin

Alternatively activated Mɸ, or those that are in a “wound healing” state are essential for myelin debris clearance following peripheral nerve injury^2,5^. We postulated that *Sarm1-/-* Mɸ at baseline or treated with IL-4 would show enhanced myelin phagocytosis compared to WT as our data suggests they are more alternatively activated. Further, our previous work showed *Sarm1-/-* mice still have lipid laden Mɸ 6 weeks post sciatic nerve injury when they should have cleared it, indicating that while phagocytosis may be intact or possibly enhanced, clearance of debris may be impaired^6^. To test this, we cultured splenic Mɸ from WT and *Sarm1-/-* mice and polarized them with either IL-4 or LPS. We then added crude myelin homogenates harvested from sciatic nerves of WT donor mice at a concentration of 1% weight/volume (**Fig3F**). Mɸ were either exposed to the myelin homogenate for 24 hours and then fixed (no clearance) or given a recovery period of 24 hours (clearance) where the myelin was removed and fresh non-myelin media was replaced (**Fig3F**). This allowed us to quantify phagocytic uptake in the no clearance, and then processing of the phagocytosed myelin in the 24 hour clearance condition (**Fig3G**). Interestingly, we found a significant increase in lipid uptake, based on OilRedO intensity in *Sarm1-/-* Mɸ treated with IL-4 compared to WT. There was no difference between genotypes at baseline or with LPS stimulation, however *Sarm1-/-* Mɸ did show a trending decrease (**Fig3G**). Additionally, when we gave the Mɸ time to process the phagocytosed myelin, we found that *Sarm1-/-* Mɸ showed overall increased OilRedO intensity compared to WT macrophages in all conditions (**Fig3G**), indicating significant decreases in clearance. In all conditions, *Sarm1- /-* mice showed overall increased OilRedO intensity compared to WT Mɸ. Altogether, this data suggests that while loss of *sarm1* does not impair myelin uptake, it does have a negative impact on clearance.

### Sarm1-/- Mɸ display mixed phenotypes following stimulation

To understand the molecular changes in *Sarm1-/-* Mɸ that could explain the outcome of the physiological assays, we turned to characterized immunophenotypes. While SARM1 is well known for its role as an NADase in axons following injury, SARM1 was originally identified as a key component of the innate immune system^16^. SARM1 is one of 5 toll-like adapter protein and suppresses signaling downstream of Toll like receptor (TLR) 4 and 2 activation through direct binding to MYD88 and TRIF proteins^16–18^. Since LPS stimulates TLR4 signaling, we posited that loss of *sarm1* should lead to enhanced downstream gene expression. Concurrently, we expected that mCSF or IL-4 treatment would have little impact on gene expression in the absence of *sarm1* since SARM1 is not known to modulate immunological stimuli receptor signaling directly. We chose several genes that are associated with either a more M1 or M2 state based on previous literature^2,5,19^. RNA was harvested from 24 hour polarized Mɸ and processed for qRT-PCR. All conditions were normalized to 12s as an internal control. Interestingly, if we compare the effect of immunological stimuli only (independent of genotype) we see differential changes in all gene expression (**FigS3**). Regardless of whether the genes are associated with more M1 or M2 immunophenotypes, *Sarm1- /-* Mɸ showed more variability between biological replicates as indicated by large standard error (**FigS3**). We transformed these data into log scale and presented it as heat maps (**FigS3B,D**). For many genes, *Sarm1-/-* Mɸ tend to increase expression of genes in both IL-4 and LPS conditions (*tnfa, nos2,tgfb, arginase, pgrn*). This was interesting because several of these genes are typically regulating opposing processes, suggesting that loss of *sarm1* dysregulates gene expression indiscriminately.

To assess the effect of *sarm1* loss on gene expression during polarization, we normalized expression to the WT mCSF condition, as this should be the most representative of homeostatic or baseline conditions. Interestingly, *Sarm1-/-* Mɸ had more dynamic changes in gene expression (although the variability was consistently higher compared to WT Mɸ), regardless of association with M1 or M2 Mɸ, and regardless of immunological stimuli (**FigS4; *arginase-1, nos2, tnfa, tgfb, il1b nmnat1, nfkb, pgrn***). Log transformation of these data and presented as heat maps indicate both stimulation dependent and genotype dependent differences in gene expression (**Fig4A,B**). In particular, *nos2* is significantly increased with mCSF and IL-4 conditions, while at the same time *arginase* is also increased in all stimulation conditions in *Sarm1-/-* Mɸ (**Fig4A,B**). Further, there is downregulation of *tnfa* in mCSF conditions, but increases in *nfkb* expression with LPS compared to WT Mɸ (**Fig4A**). These data suggest that SARM1 may be required in Mɸ for proper gene expression in response to stimuli. Without *sarm1*, Mɸ have a dysregulated gene expression that prevents the adoption of a complete immunophenotype.

**Figure 4:**
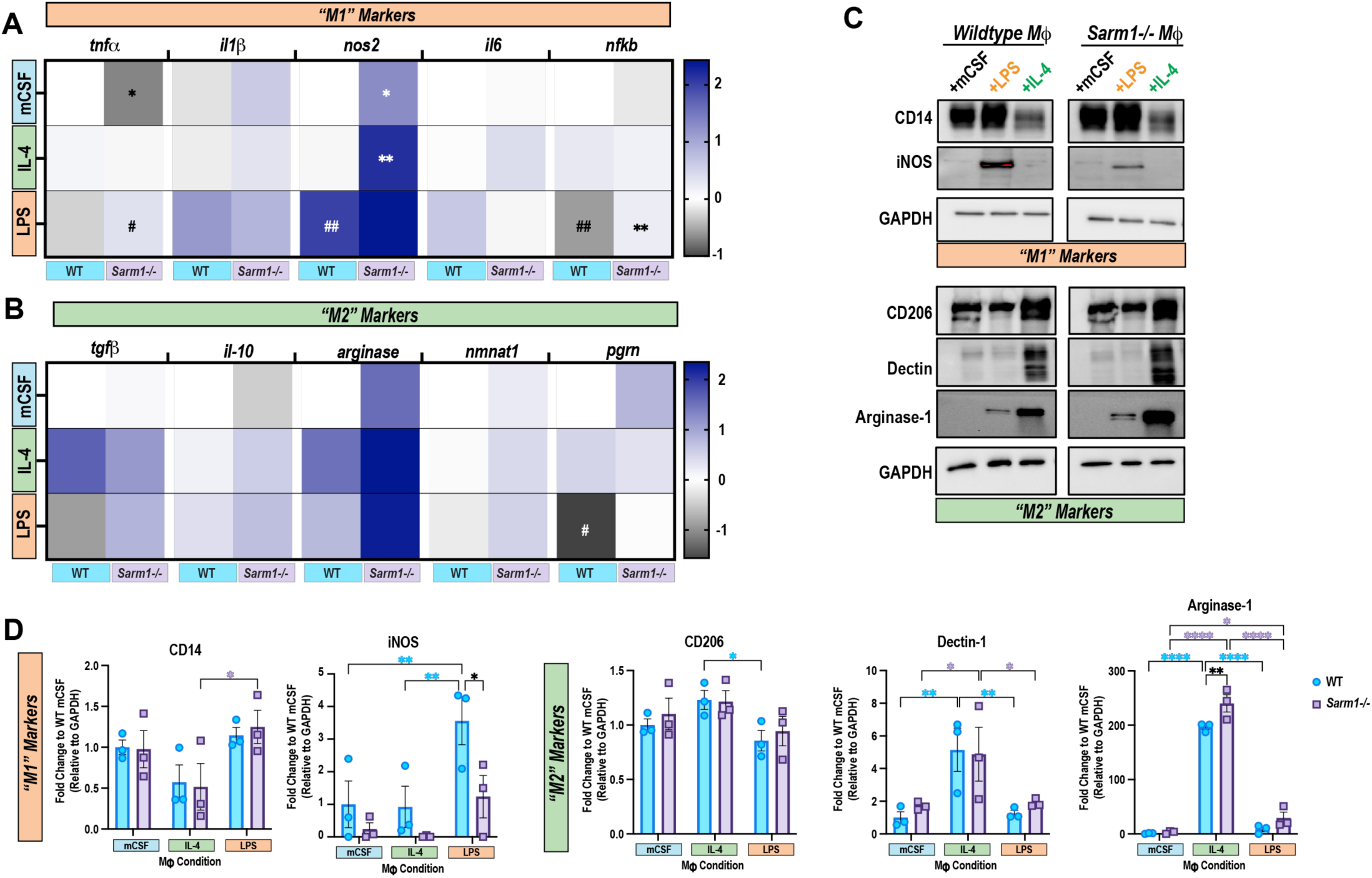
Mɸ response to immunological stimuli is partially dependent on SARM1. (**A,B**) qRT-PCR of multiple transcripts from cytokine stimulated WT and *Sarm1-/-* Mɸ. Genes are separated as indicators of M1 phenotypes (**A**) or M2 phenotypes (**B**) and represented as heat maps.12s was used as an internal control and data was represented as 2^ΔΔCt as compared to WT mCSF. Data was then log-transformed (Y=log(Y)) for visualization. N=2-4 biological replicates. *genotype comparisons; # within group comparisons. * or #p<0.05; ** or ##p<0.01 by two-way ANOVA with Fisher LSD test. (**C**) Representative western blot of stimulated WT or *Sarm1-/-* Mɸ. Blots were probed for “M1” markers (CD14, iNOS) and “M2” markers (CD206, Dectin, Arginase-1). GAPDH was used as a loading control. N= 3 biological replicates. (**D**) Quantification of relative protein levels from (**C**). Protein expression was normalized to GAPDH and then to WT mCSF and reported as a fold change. Error = SEM. *p<0.05; **p<0.01; ****p<0.0001 by two-way ANOVA with Fishers LSD test. Black asterisk = WT to *Sarm1-/-* comparisons. Purple asterisk = *Sarm1-/-* to *Sarm1-/-*comparisons. Blue asterisk = WT to WT comparisons

As another measure of immunophenotype, we probed Mɸ for proteins that are more associated with M1 or M2 phenotypes. WT Mɸ significantly upregulated M1 associated proteins like iNOS with LPS stimulation while downregulated these proteins with IL-4 stimulation (**Fig4C,D**). Similarly, WT Mɸ upregulated M2 associated proteins such as Dectin-1 and Arginase-1 with IL-4 stimulation and downregulated these proteins with LPS stimulation (**Fig4D**). Similar to our gene expression results, *Sarm1-/-* Mɸ display mixed phenotypes regardless of stimuli (**Fig4C,D**). Interestingly, there is a trending increased expression of CD14, Dectin-1 and Arginase-1 in *Sarm1-/-* compared to WT Mɸ in LPS conditions, although this is not significant (**Fig4D**). However, there is a significant upregulation of Arginase-1 in *Sarm1-/-* Mɸ with IL-4 stimulation compared to WT (**Fig4C,D)**. It is worth noting that immunological stimuli differentially impacts Mɸ within genotypes. For Dectin-1, *Sarm1-/-* show reduced up and down regulation of this protein when stimulated with IL-4 and LPS compared to WT mCSF, and levels trended towards overall upregulation (**Fig4D**). Similarly, *Sarm1-/-* failed to downregulate CD206 expression with LPS stimulation (**Fig4D**). These data suggest that *Sarm1-/-* Mɸ may overactivate M1 and M2 pathways independent of stimulation, reducing a full adoption of a singular immunophenotype.

### SARM1 is required for Mɸ immunophenotype switching

Our results suggest that *Sarm1-/-* Mɸ are more alternatively activated at baseline and treatment with IL-4 pushes them into an enhanced “wound healing” state. However, these Mɸ struggle to adopt a classical activation phenotype when challenged with LPS or IFNɣ. This led us to hypothesize that SARM1 may be required for Mɸ to switch between different immunophenotypes in response to stimuli. To explore this possibility, we designed a repolarization assay where Mɸ were initially polarized with one immunological stimuli and then after 48 hours switched to an opposing immunological stimuli (**Fig5A**). This allowed us to probe whether or not *Sarm1-/-* Mɸ could switch between “M1 and M2” states based on known molecular markers. Importantly, we not only challenged the Mɸ by switching their immunological stimuli, but we also included a replating step for some of the Mɸ (**Fig5A**). This additional challenge could either enhance or suppress the immunophenotype switch. Nine days after culturing the Mɸ we harvested lysates for western blot analysis (**Fig5A,B**). We probed for known regulators of immunophenotype switching such as Arginase-1^20^ and iNOS^21^, as well as master regulators of immunological stimuli expression, p65 and pSTAT6^22^ (**Fig5B**). Unsurprisingly, while we saw an upregulation NFKb expression and activation of p65 when Mɸ were forced to switch from an M2 to an M1 state, we did not see any differences between genotypes (**Fig5B,C**).

**Figure 5:**
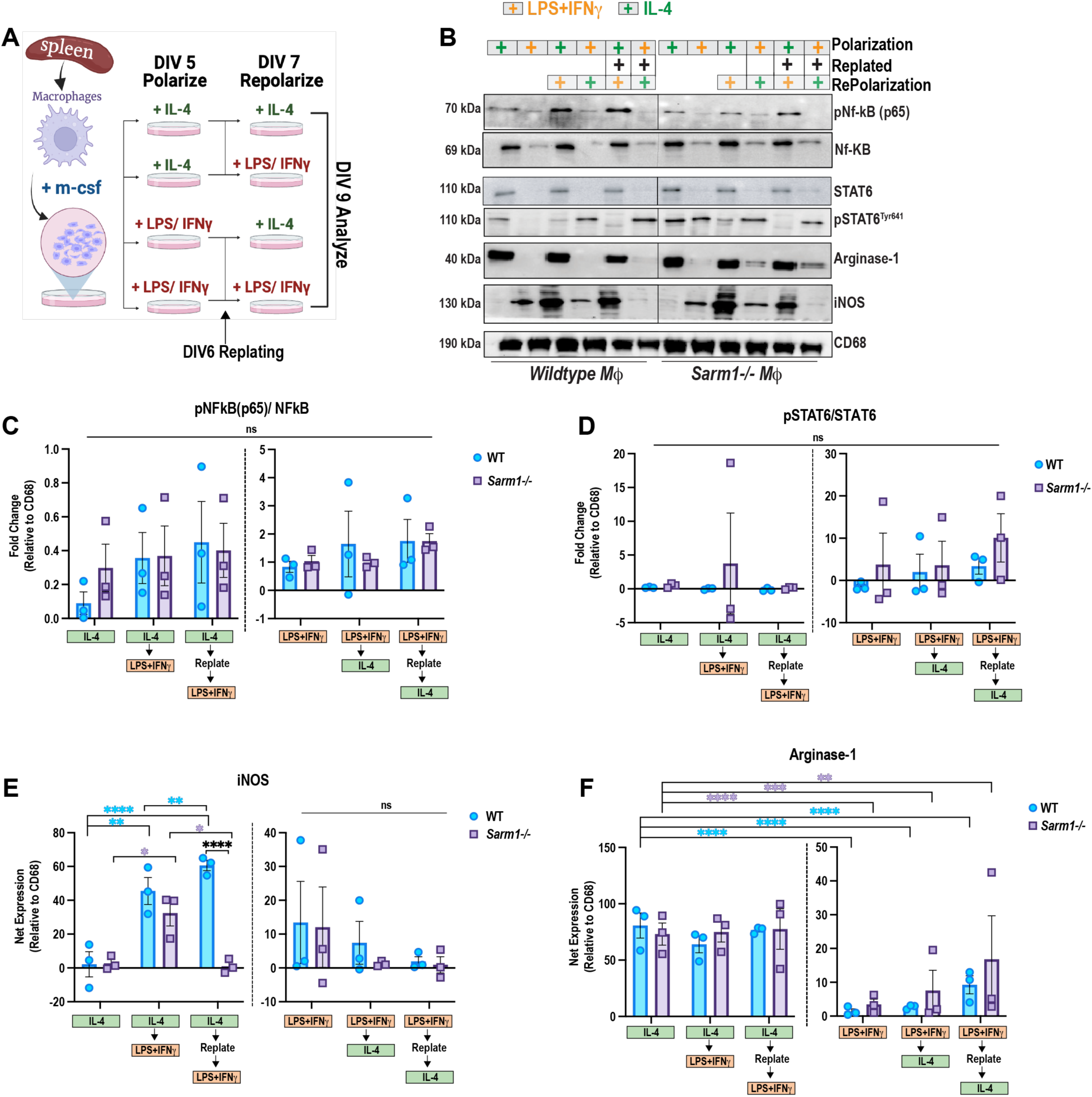
SARM1 is required for complete immunophenotype transitions in Mɸ. (**A**) Diagram of immunophenotype transition paradigm. Made in BioRender. (**B**) Western blot of WT and *Sarm1-/-*Mɸ following immunophenotype transition shown in **A**. Orange + indicates treatment with LPS/IFNɣ. Green + indicates treatment with IL-4. Black + indicates if Mɸ were replated prior to the repolarization. Samples were probed for p65, total NFkB, pSTAT6^Tyr641^, total STAT6, Arginase-1 and iNOS. CD68 was used as a loading control. (**C-F**) Quantification of protein expression from **B**. Y-axes are different between polarization conditions to improve data visualization. **C,D** represent phospho- protein levels over total levels after normalization to CD68. **E,F** represent net expression of indicated proteins compared to CD68. N = 3 biological replicates. Error bars = SEM. *p<0.05; **p<0.01; ***p<0.005; ****p<0.001 by two-way ANOVA, with Fishers LSD test. Black asterisk = WT to *Sarm1-/-* comparisons. Purple asterisk = *Sarm1-/-* to *Sarm1-/-* comparisons. Blue asterisk = WT to WT comparisons.

Previously literature has suggested that p65 activation is dependent on SARM1 expression, but this may be context specific^23^. However, we did see significant differences between genotypes in iNOS expression, and trending differences in Arginase-1 expression (**Fig5B,E,F**). Interestingly, *Sarm1-/-* Mɸ do upregulate iNOS when transitioning between M2 to M1 states, although to a lesser extent than WT (**Fig5E**). However, when these Mɸ are replated prior to the LPS addition for the repolarization, *Sarm1-/-* Mɸ fail to upregulate iNOS (**Fig5E**). This suggests that a second stimulation actually inhibits their ability to make the immunophenotype switch in response to immunological stimuli. Further, overall, *Sarm1-/-* Mɸ maintain higher levels of Arginase-1 expression regardless of immunological stimuli during the repolarization experiments (**Fig5F**). This further supports the possibility that *Sarm1-/-* Mɸ are more alternatively activated and while they are able to switch between immunophenotypes it is dysregulated and incomplete.

Since iNOS and Arginase-1 expression are linked to changes to metabolism^24^, we asked if the changes in protein expression were due to NAD consumption we performed a NAD+/NADH Glo assay in the repolarized conditions. There was no difference in overall consumption between WT and *Sarm1-/-* Mɸ, suggesting that immunophenotype switching may be independent of SARM1 NADase activity (data not shown).

### Differential neurite outgrowth from polarized Mɸ is SARM1 dependent

It has been clearly demonstrated that Mɸ are important during the neuronal injury response^2^. To determine if alterations in Mɸ phenotype from the loss of *sarm1* has other functional implications, we established a neuronal-Mɸ co-culture system. DRG neurons are easily culturable from adult rodents and have been shown to be highly sensitive to their environment^1,2,25^. Neurite length is a common measure for growth or regenerative state of the neurons *in vitro*^1,2^. To determine if the growth of neurites were impacted in the presence of Mɸ, we first plated naïve adult DRG neurons onto glass coverslips to ensure adherence of neurons in the dish. 24 hours later, we replated polarized Mɸ on top of the DRG neurons and left them in culture for another 24 hours (**Fig6A**). Using this method allowed us to assess the growth of neurites rather than their initiation which occurs during the first day *in vitro*. Neurite outgrowth was significantly enhanced in WT DRG neurons regardless of Mɸ polarization state compared to WT DRGs cultured alone (**Fig6B, D-F**). This was unsurprising as recent reports showed similar growth enhancement of DRG axons when cultured with neutrophils^26^. However, analysis within the Mɸ treatment groups showed a significant increase in both longest neurite length and branching with IL-4 treatment compared to mCSF treated controls (**Fig6D,F**). LPS treated Mɸ significantly decreased neurite length compared to IL-4 treatment but was not significantly decreased compared to mCSF treatment (**Fig6B,D**). These data suggest that DRG neurite growth is influenced by Mɸ polarization state.

**Figure 6:**
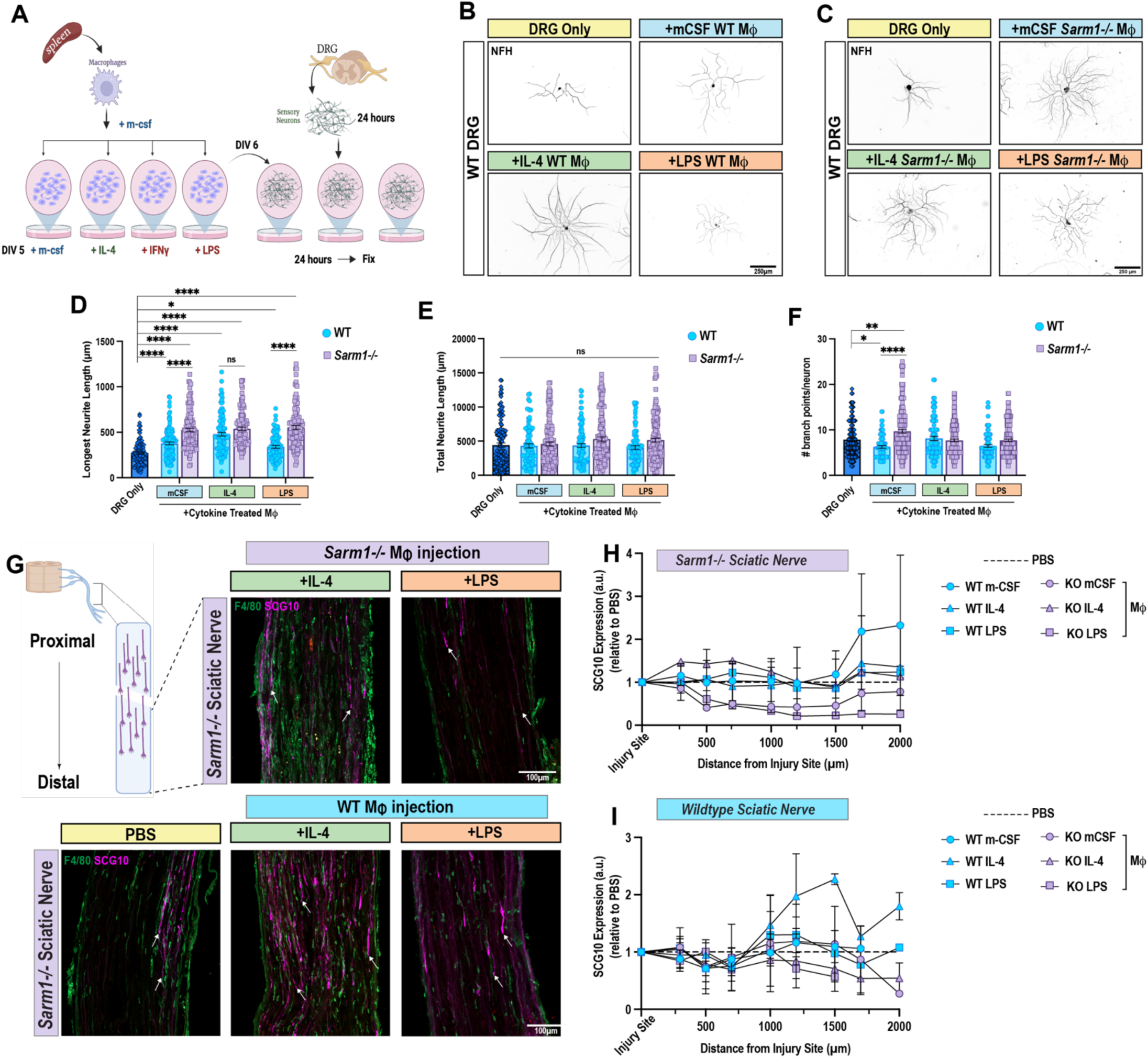
Polarized Mɸ induce changes in neurite length of sensory neurons, and loss of *sarm1* from Mɸ alters this response. (**A**) Diagram of Mɸ-sensory neuron co-culture paradigm. Made in BioRender. (**B,C**) Representative images of WT DRGs cultured alone (DRG only) or with stimulated WT (**B**) or *Sarm1-/-* (**C**) Mɸ for 24 hours (mCSF, IL-4, or LPS). Scale bar = 250µm. (**D-F**) Quantification of DRG longest neurite length (**D**), total neurite length (**E**), or number of branch points per neuron of DRGs (**F**) in **B**,**C**. Error bars = SEM (N=≥90 neurons with WT Mɸ; ≥ 88 neurons with *Sarm1-/-* Mɸ (**E**) ≥93 neurons with WT Mɸ; ≥120 neurons with *Sarm1-/-* Mɸ (**F**)≥63 neurons with WT Mɸs; ≥135 neurons with *Sarm1-/-* Mɸ from 3 independent experiments for WT Mɸ and 2 independent experiments for *Sarm1-/-* Mɸ. *p<0.05; **p<0.01; ****p<0.0001 by Kruskal-Wallis test with Dunn’s correction for multiple comparisons. (**G**) Representative images of WT or *Sarm1-/-* Mɸ injected into *Sarm1-/-* sciatic nerves. Images are 3 days after crush + injection. Mɸ are identified with F4/80 (green), and regenerating axons with SCG10 (Magenta). Arrows indicate regenerating axon tips. (**H,I**) Quantification of SCG10 expression following injection of Mɸ into *Sarm1-/-* (**H**) or WT (**I**) nerves. N=1-3 biological replicates. Expression was normalized to the injury site and represented as a fold change to the PBS control (black dashed line). Error bars = SEM.

To determine how WT Mɸ are impacting neurite growth, we performed a sholl analysis. Interestingly, there was no difference in sholl decay between all conditions (**FigS5A**). This suggests that Mɸ state does not effect neurite complexity, only total length of individual neurites. Previous reports have shown that injured PNS axons contain organized microtubule bundles^27^ and stabilization of CNS axons improves regeneration^28^. Therefore, we asked if the increase in neurite length was due to changes in microtubule stability. Here, we stained DRG neurons from our co-cultures for acetylated tubulin (**FigS5B,C**). Consistent with other reports^25^, we saw the highest expression of acetylated ⍺-tubulin in the proximal axon shaft, regardless of immunological stimuli of Mɸ (**FigS5B,C**). Suggesting that overall, stimulated Mɸ do not differentially impact microtubule stability through acetylation.

Our data suggests that polarized Mɸ differentially promote neurite length of sensory neurons. We next asked if SARM1 expression in Mɸ is required in modulating in axon growth of DRG neurons. Interestingly, *Sarm1-/-* Mɸ, regardless of immunological stimuli, also enhanced WT axon outgrowth (**Fig6C, D-F**). While there was no significant difference in neurite length between stimulation conditions, overall growth was enhanced above WT lengths when compared to stimulated WT Mɸ (**Fig6D**). Interestingly, *Sarm1-/-* Mɸ in mCSF conditions lead to significantly higher branching of WT neurons compared to WT Mɸ (**Fig6F**). These data further support a role for SARM1 in Mɸ response to stimulation. The overall enhancement of DRG neurite outgrowth regardless of *Sarm1-/-* Mɸ state, could suggest that phenotypically, *Sarm1-/-* Mɸ are more alternatively activated (anti-inflammatory) than WT Mɸ. Further, it suggests that *Sarm1-/-* Mɸ do not respond to immunological stimuli polarization to the same extent as WT Mɸ, indicating SARM1 as a potential regulator of Mɸ immunophenotype state.

Our previous work showed that *Sarm1-/-* nerves are hostile and do not support growth but grafting of WT nerves into *Sarm1-/-* can rescue the regeneration deficit^6^. Since IL-4 stimulated Mɸ enhanced outgrowth of *Sarm1-/-* DRG neurons *in vitro* we posited that we could rescue the regeneration of the *Sarm1-/-* mice by adding these Mɸ to the injured distal nerve. To do this, we polarized WT tdTomato expressing splenic Mɸ with IL-4, mCSF, or LPS as described above. We performed a sciatic nerve crush in *Sarm1-/-* mice and, immediately after and distal to the crush, we injected approximately 50,000 Mɸ. Three days later we quantified the amount of regeneration by SCG10 immunostaining. mCSF treated WT Mɸ partially rescued regeneration in *Sarm1-/-* mice compared to PBS controls (**Fig6G,H**). IL-4 treated *Sarm1-/-* Mɸ enhanced regeneration close to the injury site compared to PBS, but not at distances farther from the injury site (**Fig6H**). Both mCSF and LPS treated *Sarm1-/-* Mɸ on the other hand decreased regeneration compared to PBS (**Fig6H**). Interestingly, when WT or *Sarm1-/-* Mɸ were injected into a WT sciatic nerve, only IL-4 stimulated WT Mɸ partially rescued regeneration (**Fig6I**). This is consistent with recent reports that IL-4 itself, and IL-4 Mɸ stimulate axon regeneration^29,30^. In fact, *Sarm1-/-* Mɸ, regardless of immunological stimuli polarization, decreased regeneration in WT mice compared to PBS (**Fig6L**). These data emphasize that loss of *sarm1* impairs Mɸ response to stimulation. The inability of Mɸ to respond to environmental cues and adjust their immunophenotype likely increases the hostility of the injured nerve and does not support axon regeneration.

### SARM1 is required in both neurons and Mɸ for an efficient injury response

Our data suggest that SARM1 regulates immunophenotype switching pathways in Mɸ and the ability to undergo immunophenotype switching is important during the injury response. Injection of polarized *sarm1-/-* Mɸ, even those in an active “anti-inflammatory” state, failed to rescue regeneration *in vivo*. This suggests that immunophenotype switching is important during the injury response. To determine if loss of *sarm1* from Mɸ is responsible for the failed regeneration phenotype in germline *Sarm1-/-* animals, we generated a Mɸ conditional *sarm1* knockout line (**FigS6A**). We crossed animals that had loxP sites flanking exons 3-6 of the *sarm1* gene with a cre-line under a LysM promoter (**FigS6A**). We chose this promoter as LysM+ Mɸ are significantly upregulated in the injured nerve^31^ and we wanted to remove *sarm1* from this population of mature Mɸ. As a control, we also generated a neuronal specific *sarm1* knockout line driven with a Syn1 promoter (**FigS6A**). We first examined the state of Wallerian degeneration in these mice 7 days following SNC using the highly specific Degenotag antibody^32^ (**FigS6B**). This antibody specifically targets cleaved neurofilament light chain and does not recognize intact neurofilaments (**FigS6B**). Unsurprisingly, we saw extensive Degenotag signal in our WT mice and very little in *Sarm1-/-* mice which is consistent with our previous work^6^. Interestingly, we saw strong Degenotag signals for both the Mɸ (Mac-cKO) and neuronal (Neu-cKO) *sarm1* conditional knockout mice (**FigS6B**). Although qualitatively, the signal is reduced in the Neu-cKO which would be expected as loss of *sarm1* does delay degeneration^6^.

To further characterize the extent of Wallerian degeneration, we stained injured nerves for myelin basic protein (MBP), a phosphotidyl serine flippase (ATP8A2) which has been shown to be upregulated and a positive “eat me” signal during injury^33^, and all Mɸ (F4/80) (**Fig7A,B**). Our previous work indicated that loss of *sarm1* specifically impacts inflammation and degeneration in the distal stump, while the response at the injury site is the same as WT^6^. Therefore, we imaged and analyzed the injury site and distal stump separately (**Fig7A-C**). As expected, in WT animals at 7d post SNC we saw a reduction in MBP signal and Mɸ that appear full of ATP8A2 signals in the distal stump (**Fig7B**). In the injury site there is very little ATP8A2 signal and Mɸ have likely cleared most of the debris as we know that axons have already regenerated through the injury site at this timepoint based on previous work^6^. Consistent with our recent work, we found completely intact MBP and clear reduction in Mɸ number, and very little to no ATP8A2 signal in the distal stump (**Fig7B**). At the injury site however, the Mɸ number and morphology looks similar to WT and we can even detect the presence of ATP8A2 in the belly of the Mɸ (**Fig7A**). While the lack of ATP8A2 signal in both the WT and *Sarm1-/-* distal stump seemed perplexing, we rationalized that the axon degeneration and clearance is largely complete by 7d in the WT, but has yet to start in the *Sarm1-/-* animals.

**Figure 7:**
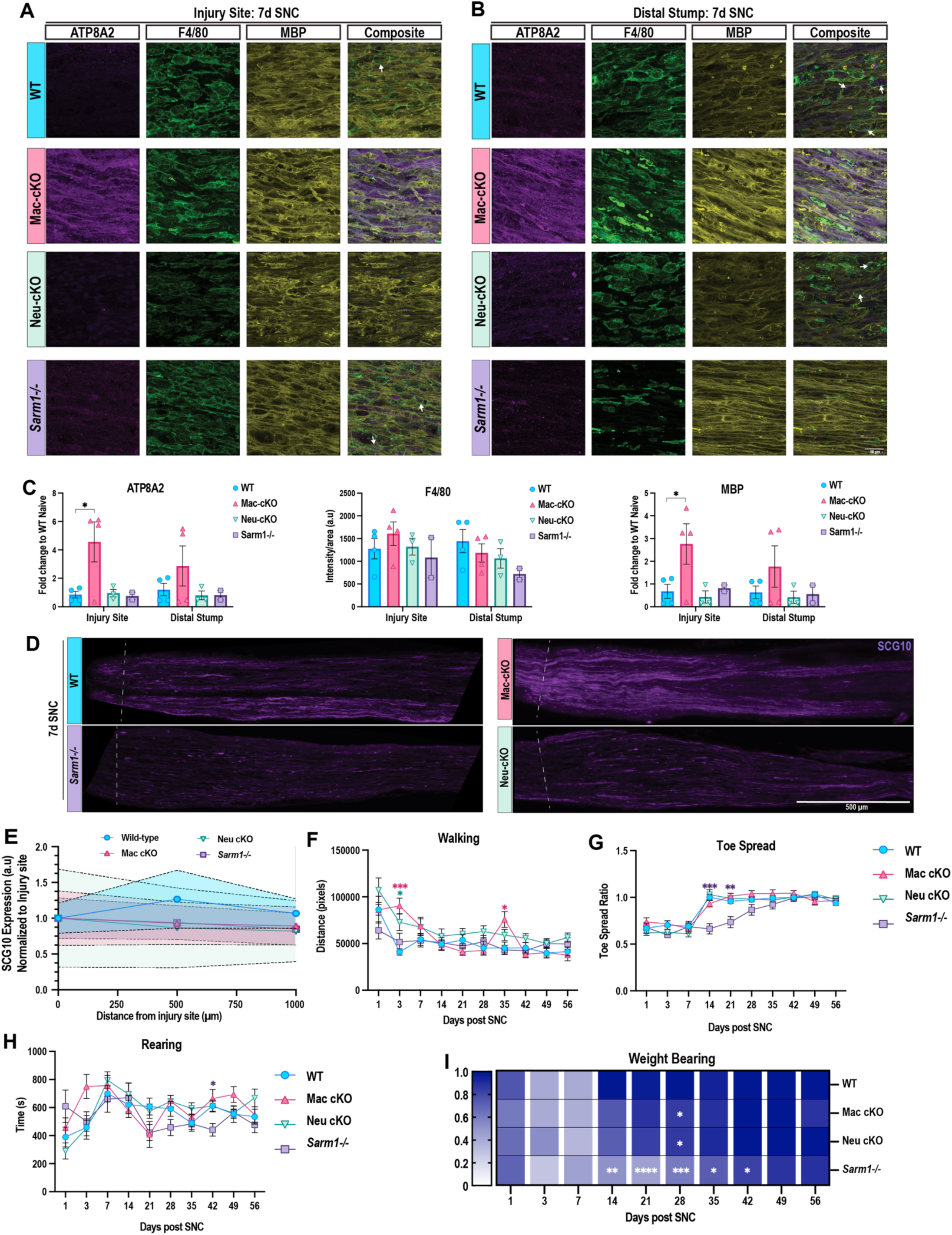
Specific loss of *sarm1* in Mɸ delays myelin clearance but does not impact overall functional recovery after SNC. (**A,B**) Representative images of injury site (**A**) and distal stump (**B**) of sciatic nerves 7 days post SNC. Phosphotidyl serine flippase (Magenta; ATP8A2), Mɸ (Green; F4/80), and myelin (Yellow; MBP). Arrows indicate F4/80 and ATP8A2 positive cells. Scale bar = 50 µm. N = 3 biological replicates. (**C**) Quantification of images in (**A,B**). ATPA82 and MBP were normalized to an uninjured WT nerve. F4/80 signal was measured as intensity/area. N=2-3 biological replicates. Data is mean +/- SEM. *p<0.05 by two-way ANOVA with Dunnett correction for multiple comparisons. (**D**) Representative images of regenerating sensory axons (SCG10; Magenta) at 7d post SNC. N=3 biological replicates. Scale = 500 µm. (**E**) Quantification of relative SCG10 intensity from (**D**) normalized to injury site. Symbols indicate mean and shaded areas indicate SEM. N= 3 biological replicates. (**F-I**) Behavior analyses from BlackBox at 10 timepoints post unilateral SNC. Data in **F-H** are represented as mean +/- SEM. (**F**) Walking distance recorded in pixels. (**G**) Toe spread ratio of injured hindlimb to uninjured hindlimb. (**H**) Time spent rearing over 20 minutes recorded in seconds. (**I**) Weight bearing ratio of injured to uninjured hind paw. 1 indicates the most pressure recorded. *p<0.05; **p<0.01; ***p<0.005; ****p<0.0001 by two-way ANOVA with Dunnett correction for multiple comparisons. Biological replicates (WT = 10; mac-cKO = 13; neu-cKO =11; *Sarm1-/-* = 12)

We then assessed the Mɸ and neuronal *sarm1* conditional knockout mice and were surprised that neither one phenocopied either WT or *Sarm1-/-* animals (**Fig7A-C**). The most striking finding was the significant signal intensity for both ATPA82 and MBP in the mac-cKO in both the injury site and distal stump compared to WT (**Fig7A-C**). Interestingly, while there was an abundance of Mɸ in present, we found very little instance of ATP8A2 signal within those Mɸ. Further, the myelin did not appear to be broken down (**Fig7A-C**). This is consistent with our *in vitro* myelin challenge assay where we saw trending decreases in myelin uptake in both mCSF and LPS conditions (**Fig3F-G**). These data suggest that loss of *sarm1* specifically in Mɸ, alters their ability to respond to injury.

Perhaps more surprisingly, we found that in the neu-cKO mice, both MBP and ATP8A2 signals were similar to WT mice in both the injury site and distal stump (**Fig7A-C**). While we expected to see a reduced ATP8A2 signal as we saw decreased axonal degeneration by Degenotag, we were surprised to see the reduction in MBP intensity and the evidence of myelin breakdown. These data suggest that the limited axonal breakdown that is independent of *sarm1*, and/or the exposed myelin fragments at the injury site is sufficient to activate Mɸ and induce clearance. This further emphasizes that Mɸ require *sarm1* to respond properly to nerve damage and argues that the delayed inflammation and degeneration in *Sarm1-/-* is secondary to loss of SARM1 activation in neurons.

To determine if myelin breakdown corresponds to sensory axon regeneration, we stained nerves 7d post SNC for SCG10 which preferentially labels regenerating sensory axons (**Fig7D**). Qualitatively, we were surprised to see that in the mac-cKO showed robust SCG10 labeling close to the injury site and within the first 500 µm distally, while the neu-cKO had very little SCG10 labeling. Quantification of the SCG10 intensity at the injury site and 500 and 1000 µm distally however showed minor reduced intensity for all *sarm1* knockout lines. However, there was variability between animals, suggesting that regeneration is likely not impaired in either the Mɸ or neuronal cKO mice (**Fig7E**).

As neither Mɸ or neuronal specific knockout of *sarm1* phenocopies *Sarm1-/-* mice, we hypothesized that neither would negatively impact functional recovery over time. To test this, we subjected male and female mice with unilateral SNC to behavior analysis using the BlackBox Bio system. Here, mice are placed in the box for 20 minute sessions at 1, 3, 7 and then weekly for a total of 8 weeks (**Fig7**). We assessed four parameters including walking distance, toe-spread, rearing, and weight bearing (**Fig7**). At 3d post SNC we found significant increase in walking distance for both mac-cKO and neu-cKO animals compared to WT (**Fig7**). Interestingly, at the same timepoint mac-cKO displayed significant increases in rearing time (**Fig7**). It is possible that the delayed response to the injury itself results in reduced functional deficits early on. Neither conditional knockout animal showed significant differences in toe spread ratio (injured foot to uinjured foot), while *Sarm1-/-* animals showed significant decreases at 14 and 21d consistent with our previous work (**Fig7**). Weight bearing showed the most differences compared to WT. Unsurprisingly, *Sarm1-/-* had a significant decrease in weight bearing on the injured foot from 14 to 42 days after SNC (**Fig7**). Both neu-cKO and mac-cKO mice showed significant decreases in weight bearing at 28d. (**Fig7**). These data suggest that expression of *sarm1* in either Mɸ or neurons is sufficient to support functional recovery after sciatic nerve injury.

## Discussion

We show that *sarm1* is partially required in Mɸ for immunophenotype transitions. *Sarm1-/-* Mɸ display both aberrant gene and protein expression during immunological stimulation and altered response to wound healing. Through DRG- Mɸ co-culture experiments, we found that Mɸ immunophenotypes differentially alter neurite outgrowth and that this is dependent on *sarm1* expression in macrophages. Finally, and consistent with our previous work, we found that *Sarm1-/-* Mɸ are hostile to the nerve microenvironment and partially prevent axon regeneration. Our data are the first to show that SARM1 expression in Mɸ is important in the context of peripheral nerve injury.

Previous studies have almost exclusively focused on the role of SARM1 in axons^7,34^. Landmark studies revealed SARM1 as a master regulator of axon degeneration as it rapidly consumes NAD+ when activated^15,35–37^. In the context of peripheral nerve injury, the distal axons undergo rapid Wallerian degeneration which is delayed in the absence of *sarm1*^6^. We previously showed, however, that loss of *sarm1* in this model leads to delayed regeneration and functional recovery, impaired immune response (primarily in the context of Mɸ), and impaired activation of repair Schwann cells^6^. Although failed Wallerian degeneration could impact immune cell infiltration and Schwann cell reprogramming via a non-cell autonomous pathway. However, recent work from the Bowie group demonstrated that bone-marrow derived Mɸ, and human monocytes, have altered glycolysis and oxidative phosphorylation when *sarm1* is removed, indicating a critical cell-autonomous role for SARM1 in Mɸ^10,11^.

Mɸ like other immune cells, rely heavily on metabolic flux to change phenotypes in response to stimulation^38^. As Mɸ transition between immunophenotypes (homeostatic, pro-inflammatory, anti-inflammatory etc.), changes in gene and protein expression lead to a shift in glycolysis or oxidative phosphorylation. Typically, Mɸ in a more “M2” or anti-inflammatory state will have increased oxidative phosphorylation and a decrease in glycolysis^38^. Two of the major upstream regulators of this shift are iNOS and Arginase-1^24^. Our data suggest, that without *sarm1* Mɸ are skewed to a more M2 state (high levels of arginase-1) and able to perform wound healing activities and phagocytose myelin debris. However, when forced to transition, or when initially polarized with pro-inflammatory cues, *Sarm1-/-* macrophages cannot fully adopt the “M1” state. Previous work has demonstrated that following sciatic nerve injury there is an influx of bone-marrow derived myeloid cells into the injury site and distal stump^2^. These infiltrating monocytes will maturate into Mɸ in the tissue over the course of the injury response^2^. In order to infiltrate the tissue these myeloid cells have to navigate from the blood and extravasate into the tissue. Our current data suggests that “M1” Mɸ without *sarm1* are unable to navigate a 3D matrix. It is therefore possible that the loss of Mɸ in the distal stump of *Sarm1-/-* mice could in part be due to failed extravasation of monocytes from the blood. This is likely due to failed regulation of cytokine production and metabolic regulation that impairs Mɸ from transitioning between immunophenotypes. We hypothesize that this impaired transition may be why *Sarm1-/-* Mɸ cannot initially support a permissive microenvironment during peripheral nerve injury, as seen in the *Sarm1-/-* mice^6^ and in our Mɸ injection model in this study.

Interestingly, we found that *Sarm1-/-* Mɸ can phagocytosis myelin debris either to the same level as WT Mɸ, or better, in the case of IL-4 stimulation. Surprisingly, there was little difference in overall OilRedO intensity in *Sarm1-/-* Mɸ between stimulation conditions, unlike their WT counterparts. Myelin clearance is a major component of the Wallerian degeneration process and is carried out primarily by Mɸ and Schwann cells^2^. Our previous work had shown that once Wallerian degeneration begins in *Sarm1-/-* mice, about 2 weeks after peripheral nerve injury, there is clear evidence of Mɸ phagocytosing debris^6^. However, interestingly these Mɸ were laden with debris 6 weeks post injury, suggesting that while they are consuming the myelin, they may not be processing it^6^. This is reminiscent of “foamy” Mɸ in the spinal cord after injury, where they are unable to process and clear lipid debris^39^. When we gave *Sarm1-/-* Mɸ an opportunity to clear the myelin, we found that they showed significant impairments compared to WT Mɸ. This was evident in control (mCSF) and IL-4 conditions, where *Sarm1-/-* displayed more OilRedO intensity in the cytoplasm compared to WT Mɸ. These data may suggest that *Sarm1-/-* Mɸ take on a “foamy” phenotype which could partly impair regeneration *in vivo*^6^.

Another consideration is the classical role of SARM1 in the Toll-Like receptor pathway (TLR) and how this may influence gene expression based on extracellular cues. While we anticipated that loss of *sarm1* would impair signaling through TLR4 activation, as SARM1 is an adapter protein downstream of TLR4, we were surprised to find that gene expression was altered regardless of immunological stimuli. It is possible that since SARM1 negatively regulates NfKb expression, there is a release of inhibition, that ultimately leads to aberrant transcriptional activation. Interestingly, we did not find increased expression of p65, but there was a trending increase in pSTAT6 which regulates many cytokine genes. It will be worth exploring if SARM1 modulates cytokine signaling in pathways other than TLR. We did note trending increases in both *nmnat1* and *prgn* gene expression which are known to promote M2 polarization. These genes would indicate other metabolic and cell signaling pathways are regulated by SARM1.

In order to dissect the cell autonomous role of *sarm1* we generated cell type specific conditional knockout animals. Much to our surprise, the neuron specific knockout did not phenocopy the regeneration and functional impairments we found in *Sarm1-/-* mice. While there was clear axonal fragmentation based on robust Degenotag staining, Mɸ were abundant and appeared to be breaking down myelin debris. Contrary to that, but consistent with our *in vitro* data, we found that Mɸ specific *sarm1* knockouts showed intact myelin signal and a decrease in Mɸ via reduced F4/80 intensity at 7 days post SNC. Intriguingly, these mice also did not show impairments in functional recovery. These data for the first time show that SARM1 expression in Mɸ is required for myelin breakdown after injury and that loss of *sarm1* from neurons alone does not block the entire injury response. Therefore, while regeneration is more efficient when Wallerian degeneration occurs, our data show that it is not entirely necessary.

## Acknowledgements

We thank the members of the SmartState Center for Childhood Neurotherapeutics for critical reading of the manuscript.

## Author contributions

Conceptualization: J.B, H.A., N.L., A.L.K; Formal analysis: J.B, H.A., N.L., A.L.K; Funding: G.B., J.S., M.S., A.L.K; Investigation: J.B., H.A., N.L., G.B., J.S., C.H., G.S., R.E.W., M.S., G.P., M.P.; Methodology: J.B, H.A., N.L., A.L.K; Project Administration: A.L.K; Resources: J.B, H.A., N.L., A.L.K; Supervision: A.L.K; Validation: J.B., H.A., N.L., G.B., J.S., C.H., G.S., R.E.W., M.S., G.P., M.P., A.L.K; Visualization: J.B., H.A., N.L., G.B., J.S., M.S., A.L.K; Writing-first draft: J.B., H.A., A.L.K; Writing- review and editing: J.B., H.A., N.L., G.B., J.S., C.H., G.S., R.E.W., M.S., G.P., M.P., A.L.K.

## Data and materials availability

*ROSA26-tdTom; Sarm1-/-*, *mac-cKO, neu-cKO,* and *Sarm1fx/fx* mice can be made available to academic researchers under a material transfer agreement upon request to A.L.K.

