## Supplemental Figures for "SARM1 is required for macrophage immunophenotype switching that is essential for nerve repair"

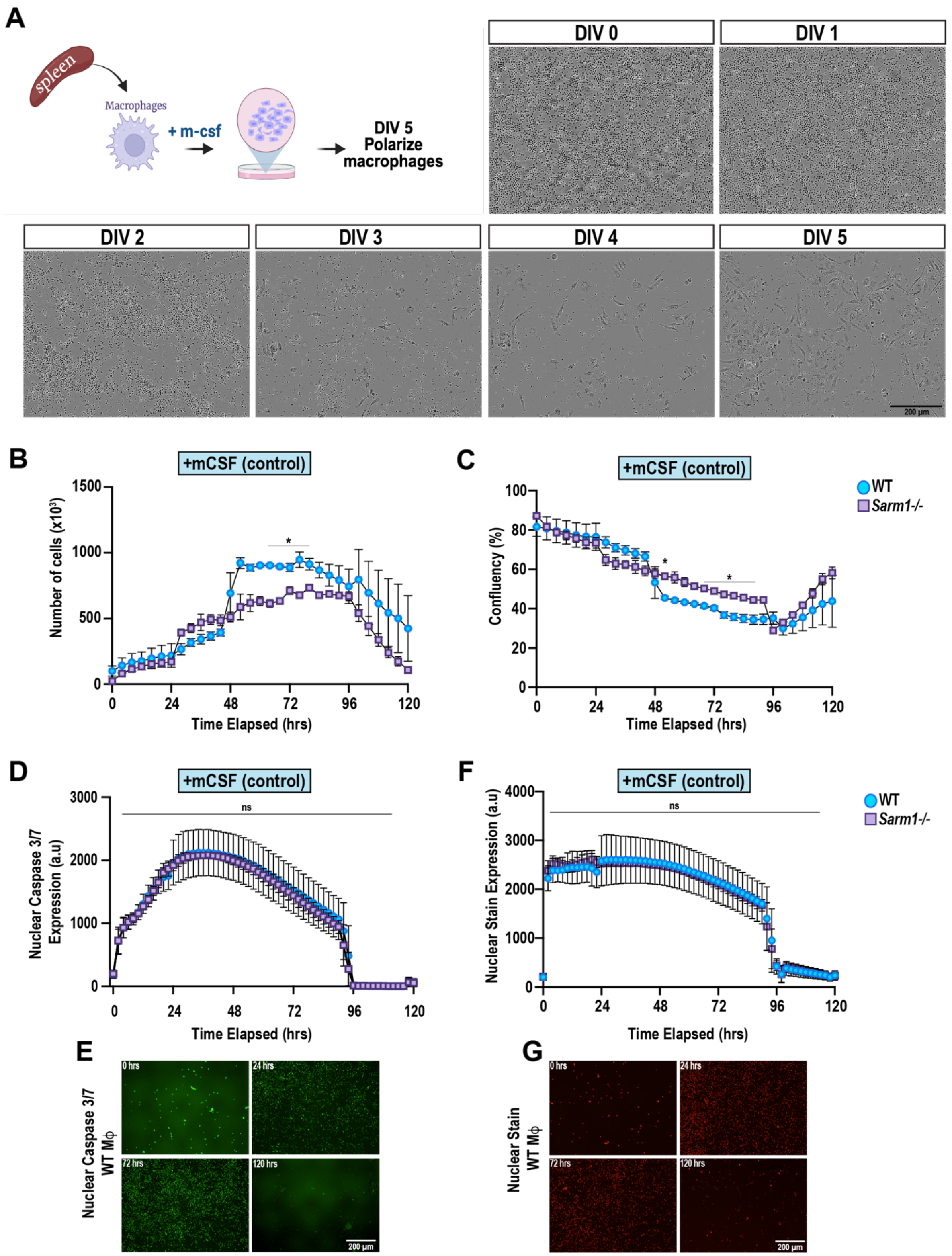


**Figure S1: Proliferation and apoptosis are not altered in *Sarm1-/-* Mϕ**

(**A**) Diagram of splenic Mϕ control culture conditions prior to immunological stimulation. Made in BioRender. Representative images from time-lapse video over the 5 day culture period. Scale bar = 200µm. (**B**) Total cell counts, (**C**) percent confluency, (**D**) nuclear caspase 3/7 expression, and (**F**) stain expression of WT and *Sarm1-/-* Mϕ during the first 120 hours in culture. N = 3 independent experiments. Error bars = SEM. *p<0.05 by two-way ANOVA, repeated measures, with Fishers LSD posthoc test. (**E,G**) Representative images from WT Mϕ cultures stained for either nuclear caspase 3/7 (**E**) or a nuclear stain (**G**) at 0, 24, 72, and 120 hrs in culture. N = 3 biological replicates. Scale bar = 200 µm.

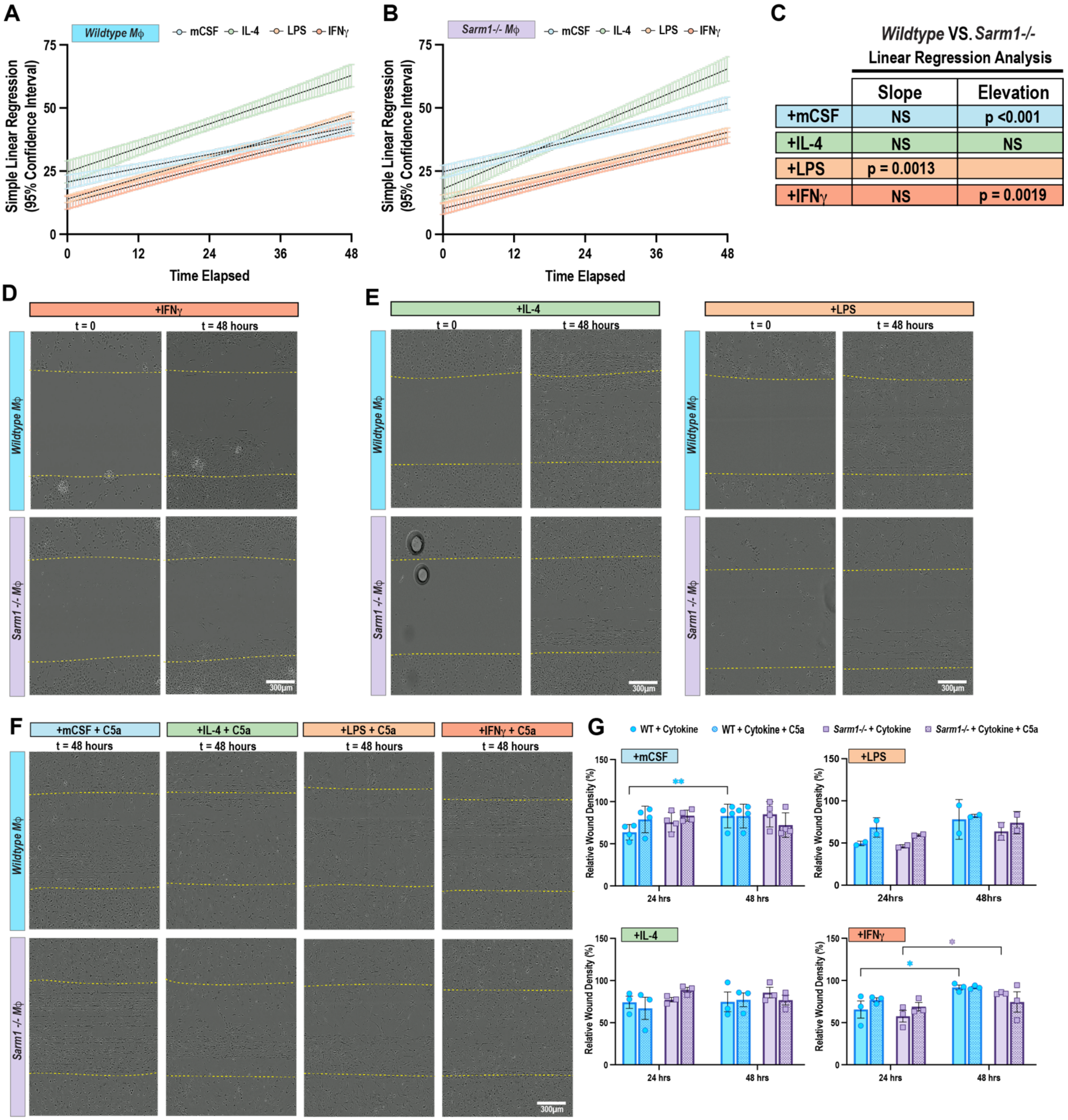


**Figure S2: Loss of *sarm1* differentially impacts Mϕ response to scratch assay**
(**A,B**) Simple linear regression of WT (**A**) and *Sarm1-/-* (**B**) Mϕ treated with mCSF (blue), IL-4 (green), LPS (light orange), IFNɣ (dark orange) after scratch assay. Error = 95% confidence intervals. N=3 biological replicates. (**C**) Simple linear regression analysis of data from **A,B**. Slope and Elevation p values are displayed. NS = not significant. Elevation for LPS could not be determined because the slope was significant. (**D-F**) Representative still images from live cell imaging sequences following a 2D scratch assay (**D**) and 3D invasion assay (**E,F**) of WT and *Sarm1-/-* Mϕ. t=0 is immediately after the scratch and t=48 hours is 48 hours post-scratch. Scratch boundaries are indicated by yellow dashed lines. Scale bar = 300μm. (**G**) Quantification of percent relative wound density at 24 and 48 hours post scratch of a 3D invasion assay. Pattern bars indicate the addition of C5a. * $p < 0.05$; ** $p < 0.01$ by two-way ANOVA with repeated measures and Tukey post-hoc test for multiple

comparisons. Blue asterisk show WT-WT comparisons. Purple asterisk show *Sarm1-/-* to *Sarm1-/-* comparisons. N = 4 biological replicates (mCSF and IL-4), 3 biological replicates (IFNɣ), and 2 biological replicates (LPS). Error bars = SEM (mCSF, IL-4, IFNɣ) and Stdev (LPS).

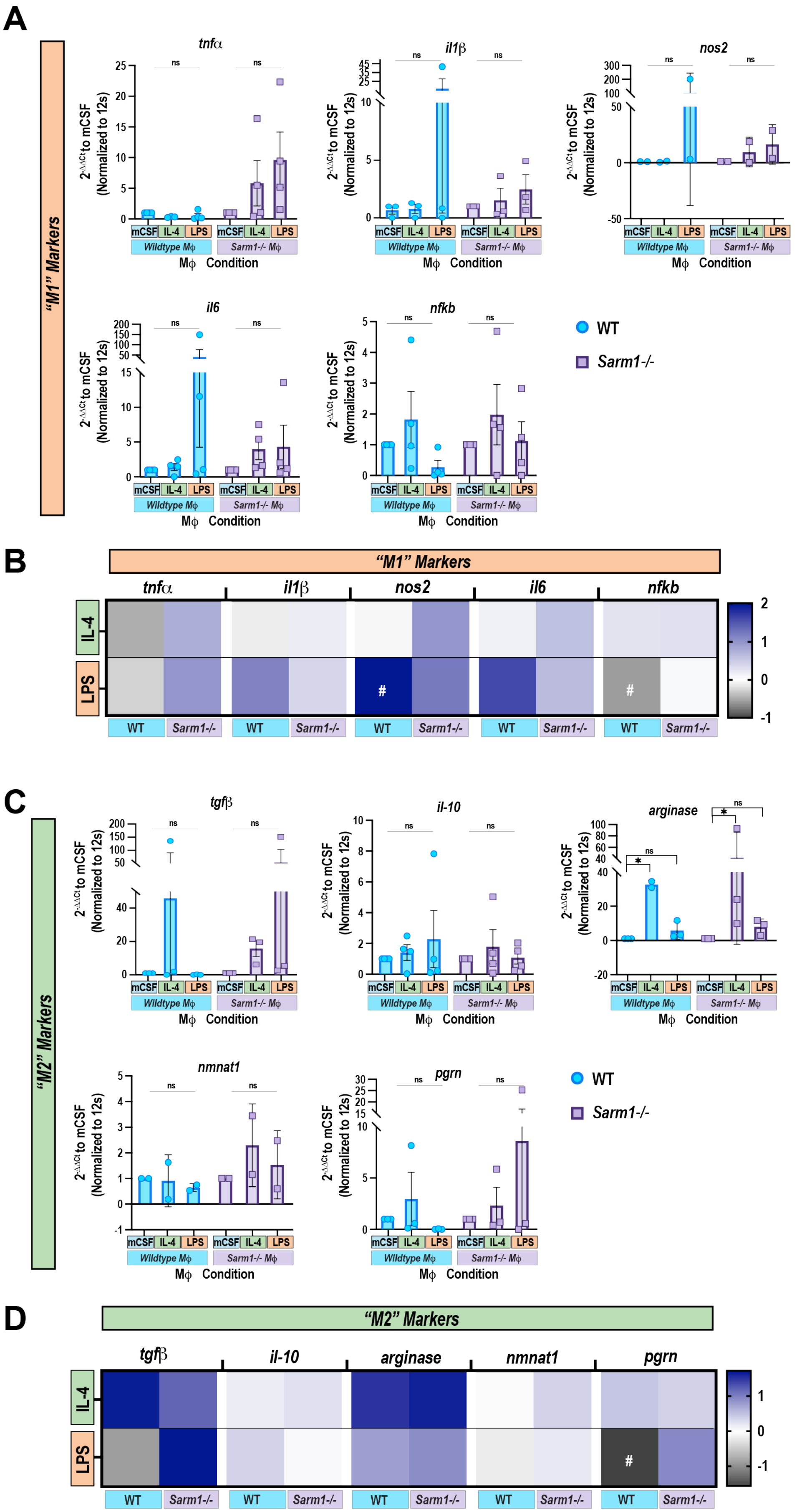


**Figure S3: Loss of *sarm1* leads to changes in stimuli dependent gene expression**

(**A-C**) qRT-PCR of multiple transcripts from cytokine stimulated WT and *Sarm1-/-* Mϕ. Genes are separated as indicators of M1 phenotypes (**A**) or M2 phenotypes (**C**). 12s was used as an internal control. Graph represents 2^ΔΔ Ct as compared to genotype specific mCSF. qRT-PCR was performed on 2-4 biological replicates and outliers were removed with ROUT Q=1%. Error is StDev for *nos2, arginase-1,* and *nmnat1*. Error is SEM for *il1b, il6, tgfb, pgrn, il-10, nfkb,* and *tnfa*. All data is non-significant by Kruskal-Wallis test and Dunn's correction for multiple comparisons. (**B,D**) qRT-PCR data from (**A,C**) represented as heat maps. Data was normalized to genotype specific mCSF and then log transformed (Y=log(Y)) for visualization. # depict within group comparisons. #$p<0.05$ by two-way ANOVA with Fisher LSD test.

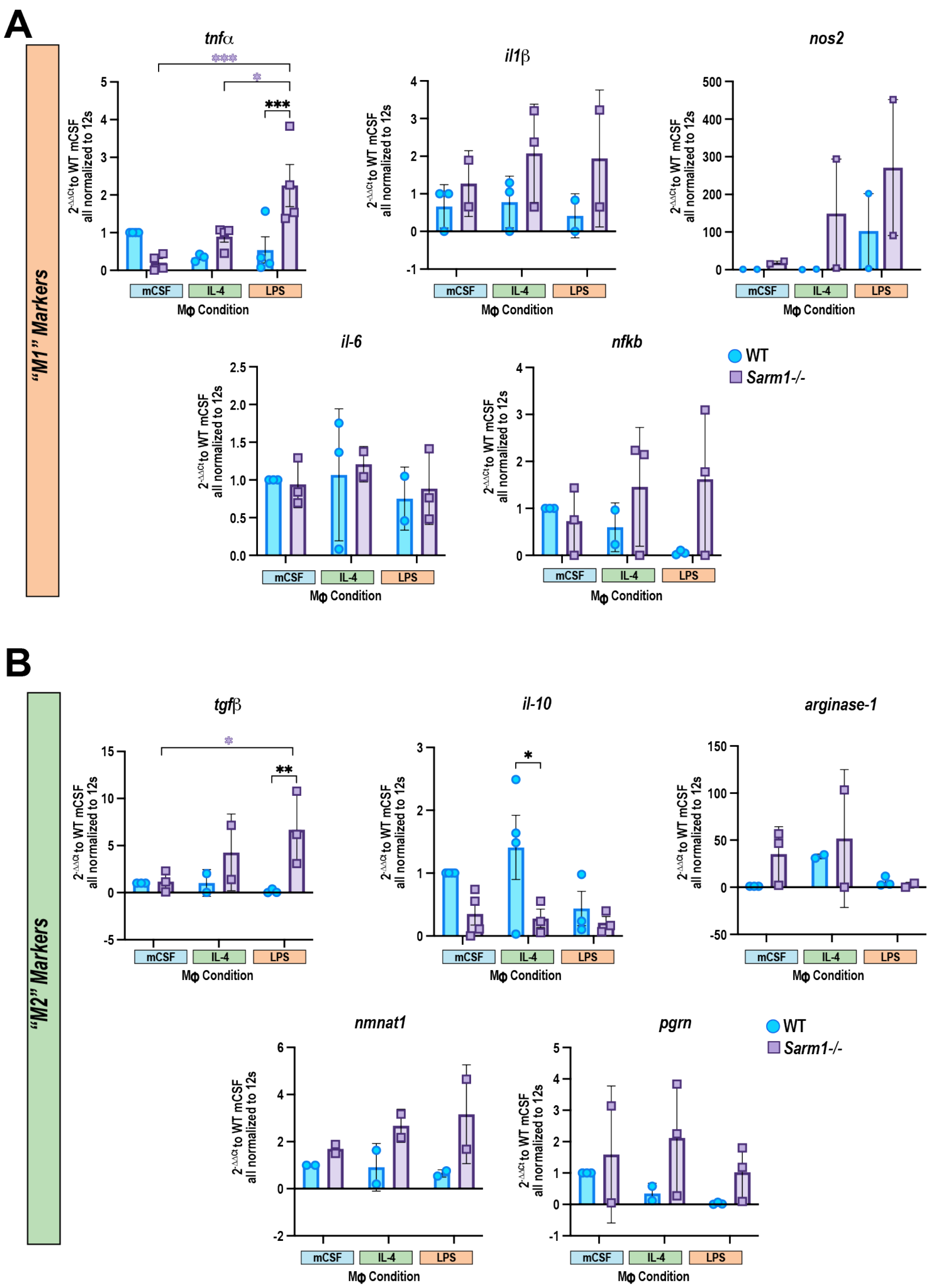


**Figure S4: Cytokine expression is dysregulated in the absence of *sarm1***

(**A**, **B**) qRT-PCR of multiple transcripts from stimulated WT and *sarm1-/-* Mϕ. Genes are separated as indicators of M1 phenotypes (**A**) or M2 phenotypes (**B**).12s was used as an internal control. Graph represents 2^ΔΔCt as compared to WT mCSF. qRT-PCR was performed on 3-4 biological replicates and outliers were removed with ROUT Q=1%. Error is StDev for *arginase-1, nos2, il1b, il-6, nmnat1,* and *nfkb*. Error is SEM for *il-10* and *tnfa*. *p<0.05; ** p <0.01; *** p<0.005 by two-way ANOVA with Tukey post-hoc test. Black asterisk = WT to *Sarm1-/-* comparisons. Purple asterisk = *Sarm1-/-* to *Sarm1-/-* comparisons. Blue asterisk = WT to WT comparisons.

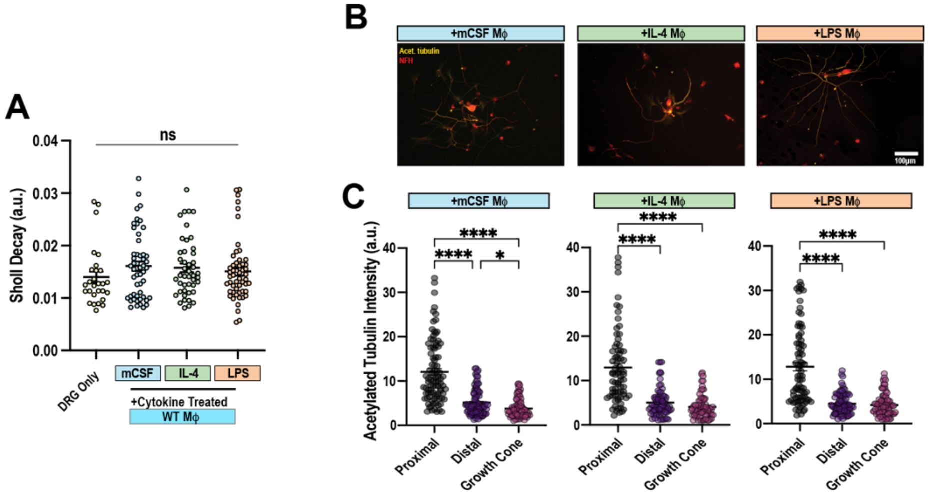


**Figure S5: Acetylation of tubulin is not impacted by polarized Mϕ**

(**A**) Sholl analysis of WT neurons with or without cytokine stimulated Mϕ. Data are represented as mean +/- SEM. N = ≥27 neurons across 3 biological replicates. (**B**) Representative images of DRGs co-cultured with either mCSF, IL-4, or LPS stimulated Mϕ. Acetylated tubulin (yellow) and NF-H (red). Scale bar = 100 µm. (**C**) Quantification of acetylated tubulin intensity in the proximal or distal axon shaft or the growth cone of WT DRGs co-cultured with either mCSF, IL-4, or LPS stimulated Mϕ. outliers were removed with ROUT Q=1%. Data is represented as mean +/- SEM. N ≥77 neurons across 3 biological replicates. *p< 0.05; **** p < 0.0001 by Kruskal- Wallis test with Dunn's correction for multiple comparisons.

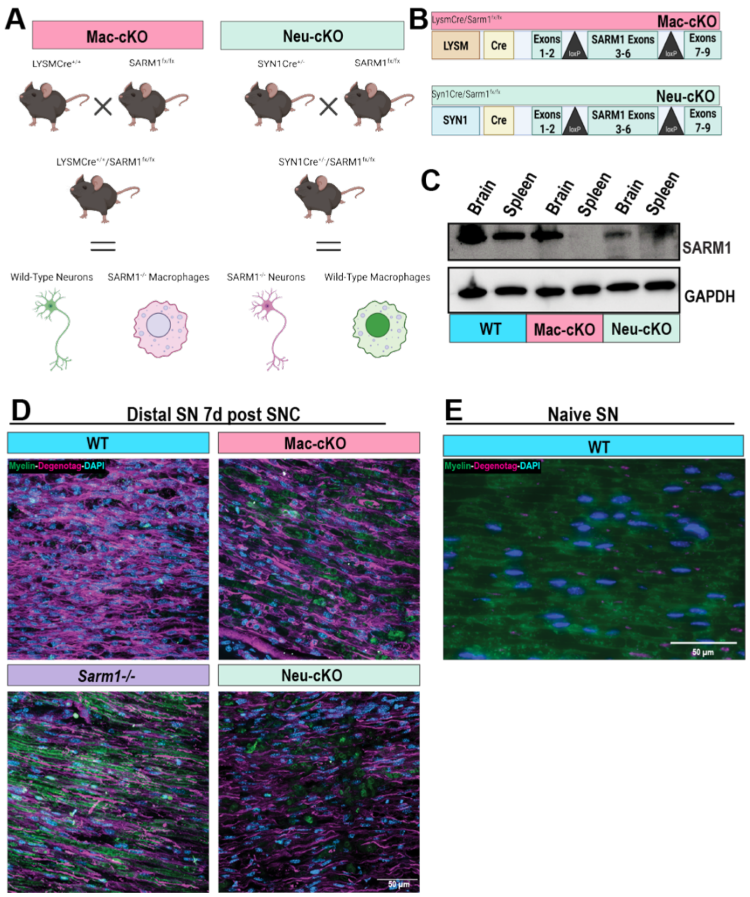


**Figure S6: Generation of conditional *sarm1* knockout mice**

(**A**) Schematic of transgenic mouse generation for Mϕ (mac-cKO) and neuronal (neu-cKO) specific *sarm1* knockout lines. Made in Biorender. (**B**) Schematic indicating where the loxP sites are in the Sarm1fx/fx mouse used for breeding in (**A**). Made in Biorender. (**C**) Representative western blot showing loss of SARM1 expression in spleens (mac-cKO) and brains (neu-cKO) in uninjured animals. GAPDH was used as loading control. (**D**, **E**) Representative images of Degenotag staining 7d after SNC (**D**) or in naïve (**E**) sciatic nerves. Myelin (Green; MBP); Degenotag (Magenta); and cell nuclei (blue; DAPI). Scale bar = 50 µm.
